# Role of *CNC1* gene in TDP-43 aggregation-induced oxidative stress-mediated cell death in *S. cerevisiae* model of ALS

**DOI:** 10.1101/2020.03.05.978411

**Authors:** Vidhya Bharathi, Amandeep Girdhar, Basant K Patel

## Abstract

TDP-43 is a multi-functional ribonucleoprotein that is also found deposited as hyper-phosphorylated and ubiquitinated TDP-43 inclusions in the brain and spinal cord of the patients of the motor neuron diseases, amyotrophic lateral sclerosis (ALS) and frontotemporal lobar degeneration (FTLD). Till date, how the cell death ensues is not fully deciphered although several molecular mechanisms of the TDP-43 toxicity such as impairments of endocytosis and chromatin remodelling, mis-regulations of autophagy and proteasome function, mis-localization to the mitochondria and generation of oxidative stress etc., have been proposed. A predominantly nuclear protein, Cyclin C, can regulate the oxidative stress response by affecting the transcription of stress response genes and also by translocation to the cytoplasm for the activation of the mitochondrial fragmentation-dependent cell death pathway. Using the well-established yeast model of TDP-43 aggregation and toxicity, we examined here whether upon TDP-43 aggregation, the cell survival depends on the presence of the *CNC1* gene that encodes Cyclin C protein or other genes that encode proteins that function in conjunction with Cyclin C, such as the *DNM1, FIS1* and *MED13* genes. We found that the TDP-43 toxicity is significantly reduced in the yeast deleted for the *CNC1* or *DNM1* genes. Importantly, the rescue of TDP-43 toxicity in these yeast deletion backgrounds required the presence of functional mitochondria. Also, the deletion of *YBH3* gene, which encodes for a protein involved in the mitochondria-dependent apoptosis, also reduced the TDP-43 toxicity. Furthermore, Cyclin C-YFP was observed to localize from the nucleus to the cytoplasm in response to the TDP-43 co-expression. Also, this cytoplasmic localization of Cyclin C was prevented by the addition of an anti-oxidant molecule, N-acetyl-cysteine. Taken together, our data suggest that Cyclin C, Dnm1 and Ybh3 proteins are important in mediating the TDP-43-induced oxidative stress-mediated cell death in the *S. cerevisiae* model.

## INTRODUCTION

Amyotrophic lateral sclerosis (ALS) is a disease characterized by the loss of motor neurons leading to atrophy of muscles, inability of movement and respiratory failure that eventually cause death [1]. Majority of the ALS cases (∼ 90-95%) are found to be sporadic (sALS), whereas 5-10% of the ALS cases are heritable & familial in nature and manifest mutations in the *TARDBP, SOD1, FUS, C9ORF72, NEK1* or several other genes [1-9]. Currently, the therapeutics of ALS remains unrealized thus uncovering the molecular mechanisms responsible for causing neuro-toxicity and cell death would be immensely helpful [2, 7, 10, 11].

The *TARDBP* gene encodes for the TAR DNA binding protein (TDP-43), which is a ribonucleoprotein with numerous roles in the RNA metabolism such as transcription, translation, stabilisation and transport of RNA etc., [7, 8]. A loss-of-functional TDP-43 pool from the nucleus due to its cytoplasmic mis-localization has been observed in the ALS and frontotemporal lobar degeneration (FTLD) disease patient’s brain and spinal cord. This has been proposed to lead to the pathological hallmarks such as the deposition of the ubiquitinated and hyper-phosphorylated TDP-43 into inclusion bodies, generation of aggregation-prone toxic C-terminal TDP-43 fragments (CTFs) and their aggregation [12-15]. In fact, this abnormal mis-localization and aggregation of TDP-43 in the cytoplasm has also been demonstrated to lead to impaired endocytosis, increased aberrant localization of TDP-43 to the mitochondria, dysregulation of the ubiquitin proteasome-mediated protein degradation, dysfunctional metal ion homeostasis, disturbances in the chromatin remodelling and oxidative stress etc. [2, 11, 16-22]. Additionally, several abnormal post-translational modifications such as cysteine oxidation, acetylation, ubiquitination, hyper-phosphorylation and PARylation have also been proposed to contribute to the TDP-43 toxicity [2, 23]. However, the final molecular events culminating into the cell death upon TDP-43 aggregation remain to be fully elucidated.

A key sub-cellular centre promising to illuminate the TDP-43 cytotoxicity is emerging to be the mitochondria. As neurons are post-mitotic cells and they also have high demands of ATP production, any mitochondrial dysfunction can severely affect their survival. Proposedly therefore, the cytotoxicity in many neurodegenerative diseases, including ALS, may be mediated *via* the mitochondrial damage [24-28]. In fact, moderate over expression of the TDP-43 has been reported to cause mitochondrial aggregation and also enhance the levels of mitochondrial fission protein Fis1 [29]. Consistent reports from several model systems, including the ALS patient-derived fibroblasts harbouring TDP-43 mutation, have also indicated of the damage to the mitochondria upon the TDP-43 expression [18, 30-34]. Notably, the overexpression of TDP-43 in the primary motor neurons alters the mitochondrial length and also impairs its transport that could be reversed upon the co-expression of the mitochondrial fusion protein, Mfn2 [18]. Also, importantly, TDP-43 and its ALS associated disease-causing mutants were found to localize to the mitochondria and preventing their abnormal localization was found to reduce the TDP-43-mediated neuronal toxicity [19, 35]. Additionally, disrupting the interaction between the mitochondrial fission proteins, Drp1 and Fis1 using a small molecule was found to mitigate the TDP-43 toxicity [29, 36]. Although mitochondrial damage and the altered expression of mitochondrial fission and fusion proteins have been consistently observed, the molecular events linked to the TDP-43-induced altered mitochondrial dynamics that initiate and culminate into cell death, remain to be elucidated. Notably, the list of human proteins that can convert *in vitro* into altered aggregation-prone amyloid-like conformation is ever increasing [37-40]. The yeast *Saccharomyces cerevisiae* has been extensively used as a eukaryotic model organism to study the aggregations and toxicities of prion and amyloid proteins of both the yeast and human origins [41-44]. Previously, expression of TDP-43 in the yeast *Saccharomyces cerevisiae* model has been shown to cause cytotoxicity and also generation of high levels of reactive oxygen species (ROS) [22, 45-48]. It is also known that mitochondria are the major centres for the generation of reactive oxygen species (ROS) and oxidative stress in a cell [49]. In addition to the mitochondrial functional dysregulation, oxidative stress has also been suggested to play vital role in the ALS pathogenesis. In fact, increasing the levels of intracellular glutathione in the motor neuron cells by supplementation with glutathione monoethyl ester, has also been shown to decrease the aggregation of TDP-43 and reduce the ROS levels [50]. Conversely, increased aggregation of TDP-43 has been observed to induce enhanced oxidative stress response through the accumulation of the anti-oxidant mediator protein, Nrf2 [27]. In addition to the mitochondria, peroxisome is another key site for the generation of reactive oxygen species as it is a dynamic organelle involved in several metabolic pathways such as fatty acid oxidation and nucleic acid catabolism [51]. Although so far, any effect of TDP-43 on peroxisome is not known, it is noteworthy that the mitochondrial fission proteins, Dnm1 and Fis1, and the fission adaptor proteins Caf4 and Mdv1, are also involved in the peroxisomal fission [52-54]. Alternatively, peroxisome fission is also regulated by Vps1 protein independently of the mitochondrial fission machinery proteins [55]. Although fission of mitochondria and peroxisomes are essential for the inheritance to the daughter cells, these fission events when not regulated can affect the yeast cell viability [53, 56].

We used the well-established yeast model of the TDP-43 aggregation and toxicity to investigate how the effect of the TDP-43 aggregation-induced oxidative stress is relayed to culminate into cytotoxicity and cell death. In yeast, and also in humans, Cyclin C which is a predominantly nuclear protein can regulate the cellular response to the oxidative stress by affecting the transcription of stress response genes and also by translocation to the cytoplasm to affect the mitochondrial fission [57]. Therefore, we examined the TDP-43 toxicity in the yeast deleted for the *CNC1* gene that encodes Cyclin C and also other genes such as *ASK10* and *MED13* encoding the proteins that function in co-ordination with Cyclin C in the event of the oxidative stress. Furthermore, we also examined the TDP-43 toxicity in the yeast deleted for the genes such as *DNM1, FIS1, CAF4* or *MDV1* that encode proteins which function in the mitochondrial fission pathway. Additionally, we also investigated the TDP-43 toxicity in the yeast strain deleted for the gene *VPS1* that codes for a regulator of the peroxisomal fission that functions independent of the mitochondrial fission response proteins. Furthermore, the TDP-43 toxicity was also examined in the yeast strain deleted for the *YBH3* gene that encodes for a pro-apoptotic protein that gets activated in response to the mitochondrial hyper-fission. Finally, we investigated if the oxidative stress caused by the TDP-43 protein aggregation can alter the sub-cellular localization of the Cyclin C protein and whether neutralizing the oxidative stress by the addition of an anti-oxidant can thwart its cytoplasmic localization induced by the TDP-43 co-expression.

## MATERIALS AND METHODS

### Materials

D-raffinose, 2, 3, 5-triphenyl tetrazolium chloride, D-galactose, ampicillin, SDS, 2’,7’ – dichlorofluorescin diacetate (DCFDA) and lithium acetate were purchased from Sigma (USA). D-glucose was procured from Amresco (USA). Yeast nitrogen base, yeast extract, peptone, DMSO, chloroform, agar and glass slides were procured from HiMedia (India). Salmon sperm DNA was purchased from Invitrogen (USA). CellRox Deep Red reagent (catalogue no: C10422) was purchased from Thermofisher scientific. For western blot, CellLytic™ Y (Sigma USA), Bradford reagent (BioRad), Western blot blocking buffer (Himedia India), anti-TDP-43 primary antibody (Sigma, catalogue no: WH0023435M1), anti-GAPDH primary antibody (Abcam, catalogue no: ab125247), anti-mouse secondary antibody tagged with alkaline phosphatase (Sigma, catalogue no: A3562), CDP star (sigma), and molecular weight standards (Thermofisher scientific), were used. Propidium iodide (Sigma, catalogue no: P4170) was used for dead cell staining. For staining the yeast nuclei, DAPI (Sigma, catalogue no: D9542) or Hoechst 33342 (Thermofisher scientific, catalogue no: 62249) were used.

### Yeast strains

Yeast strains BY4742 (*MAT****α*** *his3*Δ*1 leu2*Δ*0 lys2*Δ*0 ura3*Δ*0*) (obtained from GE Dharmacon) and 74-D-694 (*MAT****a*** *ade1-14 his3-200 ura3-52 leu2-3, 112 trp1-289*) were used for the studies [58]. The other yeast strains used including *ask10Δ, med13Δ, fis1Δ, dnm1Δ, ybh3Δ, mdv1Δ, caf4Δ, vps1Δ* and *cnc1Δ*, were all derivatives of BY4742 and obtained from the yeast *MAT alpha* knock out clone collection from GE Dharmacon [59, 60]. Isogenic [*pin*^-^] versions of the *ask10Δ, med13Δ, fis1Δ, dnm1Δ, ybh3Δ, mdv1Δ, caf4Δ, vps1Δ* and *cnc1Δ* yeast strains were obtained by growing the strains on YPD plates supplemented with 5 mM GuHCl for five passages as patches to eliminate any presence of the [*PIN*^*+*^] prion. All studies were carried out in the [*pin*^*-*^] yeast strains [61].

### Plasmid construction

Plasmids used in the study were *pRS416 (URA3)[62]*, low copy Tdp-43 expression plasmid *pRS416-GAL1p-TDP43-YFP (URA3)* (Addgene: 27447), high copy Tdp-43 expression plasmid *pRS426-GAL1p-TDP43-GFP (URA3)* (Addgene: 27467), *pGAL1-TDP-43-DsRed (URA3)* and *pGAL1-TDP-43 (TRP1)* [45, 46, 63]. Also, a plasmid expressing a C-terminal fragment of TDP-43 (aa: 175-414) as GFP fusion was generated here using Q5 site-directed mutagenesis system (NEB, USA) using the plasmid *pRS426-GAL1p-TDP43-GFP* (Addgene: 27467) as the template following the manufacturer’s protocol. The forward primer 5’*ATGTGCAAACTTCCTAATTCTAAGC*3’was used along with the reverse primer 5’*TTTGTTATCTCCTTCGAAGCCTG*3’ to generate the plasmid (*pRS426-GAL1p-TDP-43-(175-414)-GFP*) to express the C-terminal fragment aa: 175-414 of TDP-43 tagged with GFP. For the plasmid construction, following the *DpnI* digestion of the template DNA, the PCR product was transformed into competent *DH5α E. coli* cells (Invitrogen, USA). Then, plasmids were isolated from the obtained transformants and analyzed for their respective desired lengths of DNA by double digestion using restriction enzymes *SpeI* (which cuts before initiation codon of TDP-43 encoding gene) and *XhoI* (which cuts after *GFP* gene) to ascertain for the successful generation of the C-terminal fragment encoding plasmid. In addition, to generate a Cyclin-YFP encoding plasmid, a *CNC1* ORF obtained from GE Dharmacon (catalogue no: YSC3867-202329183) was used as the template. The forward primer 5’*ACTTCTAGAATGTCGGGGAGCTTCTGG*3’was used along with the reverse primer 5’*ATGGATCCAATTGCAGATGCTGGTCTAAG*3’ to PCR amplify the *CNC1* ORF added with *XbaI* and *BamHI* restriction sites in the forward and the reverse primers respectively. The plasmid *pRS426-GAL1p-RNQ1-YFP* (Addgene: 18686) was first digested with the restriction enzymes *XbaI* (which cuts before initiation codon of the *RNQ1* gene) and *BamHI* (which cuts before the *YFP* gene). Then the *RNQ1* ORF insert was removed and the vector with *YFP* was used to ligate in the *XbaI* and *BamHI* digested PCR product of the *CNC1* ORF to obtain the *pRS426-GAL1p-CNC1-YFP* plasmid.

### Yeast culture, plasmid transformation and diploid generation

Standard procedures for the yeast media preparation and cultivation were used [64]. The yeast cells were grown either in the complex media (YPD) or in a standard synthetic media lacking a particular amino acid and containing either D-glucose (SD) or D-raffinose (SRaf) or galactose (SGal) as the sole carbon source. Transformation of yeast cells with plasmids was carried out using the standard Li-Acetate method [64]. Galactose (2%) was used to over-express the *GAL1* promoter-controlled genes from the plasmids. For generation of the diploid strains towards complementation assays, a yeast strain deleted for a specific gene in the BY4742 background (*MAT****α***) was mated with *74-D-694* wild-type strain (*MAT****a***) on a YPD plate and then the diploids were selected on SD-Lys-Trp plates [58].

### Yeast growth inhibition assay

Yeast cells were first cultured overnight in plasmid selective, 2% raffinose containing broth to overcome the glucose repression. Next morning, the cells were inoculated to a fresh raffinose containing broth and allowed to grow till mid-log phase OD_600nm_=0.5. Then, the cell numbers were normalized and cells were serially diluted by ten-folds and 5 μL of the cell suspension was spotted onto appropriate plasmid selective synthetic media plates that contained glucose, raffinose or galactose as the sole carbon source. The plates were incubated at 30 °C for 2 days before imaging. Any toxicity due to the expression of TDP-43 through the galactose promoter was assessed by comparing the relative growths on the galactose containing plates in reference to appropriate controls.

### Generation of petite yeast strains

Isogeneic petite [*rho-*]versions of *dnm1Δ, ybh3Δ* and *cnc1Δ* were created by growing the [*RHO*^+^] strains on YPD supplemented with 0.5 mg/ml ethidium bromide (EtBr) which has been previously reported to result in respiratory-deficient mutant strains termed [*rho-*] or petite strain [65]. Cells from the EtBr plates were then streaked on to YPD plates to obtain single colonies. Several colonies from the YPD plates were analysed for screening the respiration-deficient mutants (i.e. conversion to [*rho-*]) by checking for their lack of growth on the complex media YPG containing 2% glycerol (non-fermentable) as the only carbon source.

### Western blotting

To examine the protein expression levels of TDP-43-YFP or TDP-43-GFP, the cell extracts from the yeast cells grown for 24 h with 2% galactose were analyzed by western blotting. Yeast cells were lysed by vortexing at 4 °C with glass beads in CellLytic™ Y buffer and the lysate was pre-cleared of the cellular debris by centrifugation at 2000 rpm at 4 °C for 3 min in a table-top micro-centrifuge. The supernatant was decanted out and the total protein concentration was estimated using Bradford assay. Normalized amounts of samples with equal total protein contents were electrophoresed on 10% SDS-PAGE and electro-blotted to polyvinylidene difluoride (PVDF) membrane. After this, the membrane was incubated for 120 min with casein blocking buffer (HiMedia) and washed with phosphate buffered saline (PBS) containing 0.1% Tween (pH 7.5). Mouse monoclonal anti-TDP-43 primary antibody (1:2000 dilution) was used for probing TDP-43-YFP and TDP-43-GFP followed by an incubation with anti-mouse secondary antibody (1:10000 dilution) tagged with alkaline phosphatase (APP). Protein levels of the endogenously expressed GAPDH were also detected concurrently as the protein loading controls. For this, the mouse monoclonal anti-GAPDH primary antibody (1:10000 dilution) tagged with APP was used. CDP-star was used as the chemi-luminescent substrate for APP as per the manufacturers protocol to visualize the levels of the TDP-43 and GAPDH proteins. A molecular weight standard cocktail was electrophoresed and electro-blotted simultaneously for assignment of the molecular weights to the TDP-43 and GAPDH proteins.

### Fluorescence microscopy

For fluorescence microscopy, single colony isolates of the yeast plasmid transformants were grown overnight at 30 °C in raffinose containing broth media. Then, the cells were re-suspended in fresh media and expressions of TDP-43-YFP, TDP-43-GFP or Cyclin C-YFP from the *GAL1* promoter were induced with 2% galactose and incubated at 30 °C before harvesting the cells for microscopy. Fluorescence was observed under Leica DM-2500 microscope and images were acquired using 100x objective lens (oil immersion) in bright field or by using the YFP or GFP filters and further processed using ImageJ software [66]. Detection of presence of oxidative stress in yeast upon TDP-43-YFP expression using CellROX deep red staining and visualization of staining of yeast nuclei by DAPI or Hoechst were also analysed by fluorescence microscopy as detailed in the following sections.

### CellROX deep red staining for ROS detection

Presence of reactive oxygen species (ROS) was detected by staining the yeast cells with CellROX deep red dye followed by visualization of the cells in microscope using RFP filter. For this, the yeast cell suspensions expressing *URA3* marked plasmid encoded TDP-43-YFP or vector were prepared as described earlier [67]. First, yeast cultures were grown overnight in SRaf-Ura and inoculated into fresh SGal-Ura for the overexpression of the TDP-43-YFP for 24 h. Then, the cells were pelleted and re-suspended in 100 μL of water and incubated for 1 h with 1.5 μM CellROX deep red in dark. Fluorescence was detected under Leica DM-2500 microscope and cell images were acquired using 100x objective lens (oil immersion) in bright field and also RFP filter and then processed using ImageJ software [66].

### DAPI and Hoechst staining for nuclei visualization

To examine the cytoplasmic localization of TDP-43-YFP aggregates, after 24 h of expression of TDP-43-YFP, the yeast cells were collected by centrifugation and re-suspended in 70% ethanol and incubated for 30 min at room temperature. Then, the cells were washed and re-suspended in PBS and incubated with 5 μg/mL of DAPI for 10 min at room temperature. After incubation, the yeast cells were washed thrice with PBS before visualizing the nuclei under microscope. For examining the nuclear localization of Cyclin C-YFP, the nuclei were stained with the Hoechst dye. For this, Cyclin C-YFP was expressed either alone or co-expressed with TDP-43-DsRed for 18 h before the yeast cells were collected by centrifugation. Next, the cells were washed with PBS and incubated with 5 μg/mL of Hoechst for 30 min at room temperature. After incubation, the cells were washed thrice with PBS before visualizing the nuclei under Leica DM-2500 microscope.

### Flow cytometry

Yeast cells were induced using 2% galactose for 36 h to express TDP43-YFP or EGFP. More than one million cells were harvested and washed with PBS. Next, the yeast cells were re-suspended in 500 μL of PBS and stained with 10 μg/mL of propidium iodide (PI) and incubated for 15 min in the dark. Then, 50,000 cells were acquired in AriaIII FACS (BD) and analysed using the filter PI-A (for PI) to calculate the percentage of total dead cell population which get stained with PI. For examining the oxidative stress levels in the yeast, CellROX deep red stained population was also quantified by flow cytometry. For this, the yeast cells harbouring TDP-43-YFP or EGFP encoding *GAL1* promoter-driven plasmid, were induced using 2% galactose for 36 h and then harvested and washed with PBS. Next, the yeast cells were re-suspended in 500 μL of PBS and stained with 10 μM of CellROX deep red reagent for 1 h in the dark. Then, 50,000 cells were acquired in AriaIII FACS (BD) and analysed using the filter APC-A (for CellROX) to calculate the percentage of CellROX-stained cells in a given population.

## RESULTS

### Deletion of *CNC1* and some associated mitochondrial pathway genes rescue the TDP-43 toxicity

Several mechanisms of the TDP-43 aggregation-induced cytotoxicity have been proposed thus far such as the interferences with the RNA metabolism, proteasome function, mitochondrial function, endocytosis and lysosome function etc. [20, 22, 63, 68, 69]. TDP-43 expression has also been shown to induce mitochondrial damage, display elevated production of reactive oxygen species (ROS) and impair the mitochondrial membrane potential [35]. In fact, TDP-43 has been shown to localize to the mitochondria and the inhibition of its mitochondrial localization can significantly reduce the neuronal toxicity [19]. Also, TDP-43 has been found to interact with certain mitochondrial proteins that are important for mitophagy [70]. Although mitochondrial dysfunction has been observed with TDP-43 expression, the molecular mechanism by which TDP-43-induced mitochondrial damage culminates into cell death is not well understood. In yeast, the Cyclin C protein can mediate the stress response against various stresses including the oxidative stress, heat shock, osmotic shock, actin depolymerisation and cell wall stress [71-74]. Of note, Cyclin C along with its partner Cyclin dependent kinase (Cdk8) associates with the RNA polymerase II and represses stress response genes **(Figure 1)**. In the event of oxidative stress, Cyclin C dissociates from Cdk8 as well as its nuclear anchor protein Med13 and translocates to the cytoplasm where it associates with the mitochondria [75]. Ask10 is another important component of the oxidative stress signal pathway which affects the Cyclin C’s dissociation from Cdk8 and Med13 to translocate into the cytoplasm [76]. Upon its cytoplasmic localization, Cyclin C associates with the mitochondria and helps in signalling the recruitment of the mitochondrial fission machinery [75]. The mitochondrial outer membrane receptor, Fis1, recruits the dynamin-like GTPase protein, Dnm1, to the mitochondria *via* two adaptor proteins, Mdv1 and Caf4 [77]. This helps in activating the enhanced mitochondrial fission response which in turn relays into the movement of the pro-apoptotic protein, Ybh3, from the vacuole to the mitochondria that can cause programmed cell death (PCD) (**Figure 1**) [57]. As TDP-43 expression causes oxidative stress in the yeast cells, we examined if Cyclin C and its associated proteins are involved in mediating the TDP-43-induced cell death.

**Figure 1:**
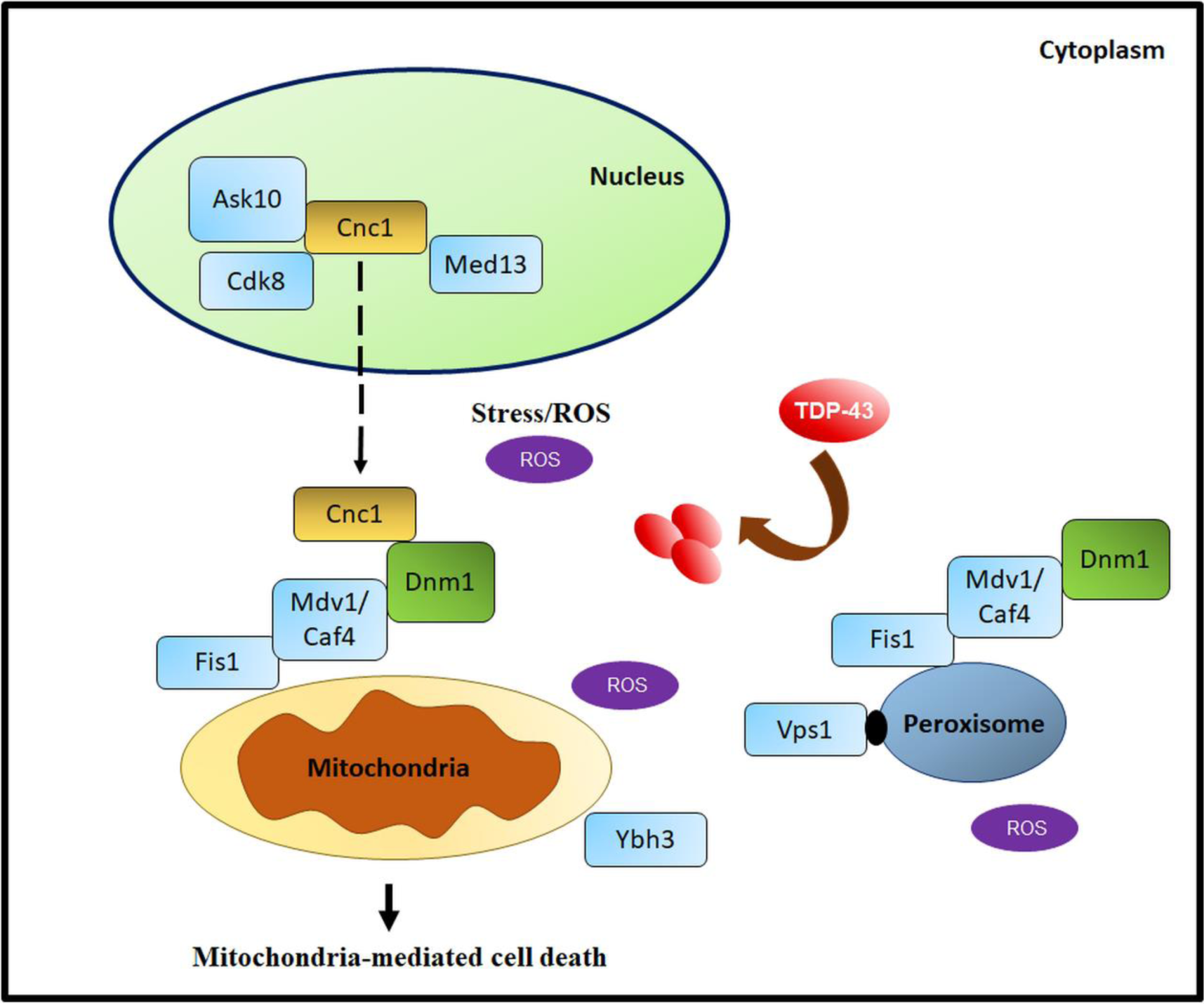
Schematic of cytoplasmic localization of Cyclin C upon oxidative stress and protein complexes that function in the fission of mitochondria & peroxisomes in yeast. Mitochondria and peroxisomes are the sources of reactive oxygen species (ROS) and their fission can regulate cell death and longevity. Oxidative stress-induced cell death in yeast is mediated *via* mitochondria. Various stress signals, including oxidative stress, elicit Cyclin C protein-mediated stress response in yeast. Cyclin C protein along with the Cyclin dependent kinase, CDK8, is a predominantly a nuclear protein. Upon oxidative stress, Cyclin C dissociates from CDK8 as well as from its nuclear anchoring protein Med13 and moves from the nucleus to the cytoplasm. Ask10 protein activation in response to the oxidative stress also promotes the Cyclin C’s movement to the cytoplasm. After localizing to the cytoplasm, Cyclin C triggers the recruitment of the dynamin-like protein, Dnm1, to the mitochondrial fission protein Fis1 *via* the adaptor proteins Caf4 or Mdv1. This activates mitochondrial fission. Aberrant mitochondrial fission also leads to the recruitment of the pro-apoptotic protein, Ybh3, from the yeast vacuole to the mitochondria which thereafter starts the programmed cell death (PCD) pathway. The mitochondrial fission machinery proteins Fis1, Caf4 and Mdv1 are also involved in regulating the peroxisomal fission in a Dnm1-dependent manner. Another protein, Vps1, also regulates the peroxisomal fission in a Dnm1-independent manner. We examined if the TDP-43 aggregation-induced oxidative stress can also affect the yeast cell death *via* the Cyclin C-mediated mitochondria-dependent PCD pathway.

First, we performed a serial dilution growth assay in the wild-type yeast strain transformed with a low copy plasmid encoding TDP-43-YFP driven by the *GAL1* promoter and found that the expression of TDP-43-YFP caused cytotoxicity as reported earlier (**Figure 2a**). As TDP-43-YFP expression is also known to cause oxidative stress in yeast, we therefore examined the TDP-43-YFP toxicity in the yeast strains deleted for the Cyclin C encoding gene *CNC1* or the downstream genes that encode for the proteins that work in collaboration with Cyclin C. Strikingly, upon TDP-43-YFP expression, the deletion of *CNC1* and also that of the *ASK10* gene whose encoded protein is essential for the translocation of Cyclin C from nucleus to the cytoplasm, exhibited rescue of the TDP-43-YFP toxicity, whereas, the deletion of the *MED13* gene encoding the Med13 protein which is essential for retaining the Cyclin C in the nucleus, did not exhibit any rescue of the TDP-43 toxicity (**Figure 2a**). Furthermore, we also examined the toxicity of TDP-43-YFP in the yeast deleted for the genes encoding the proteins Dnm1 and Fis1 that are involved in the mitochondrial fission process. We found that the TDP-43-YFP toxicity was reduced in the d*nm1Δ* yeast strain, whereas, there was no toxicity rescue in the *fis1Δ* yeast (**Figure 2a**).

**Figure 2:**
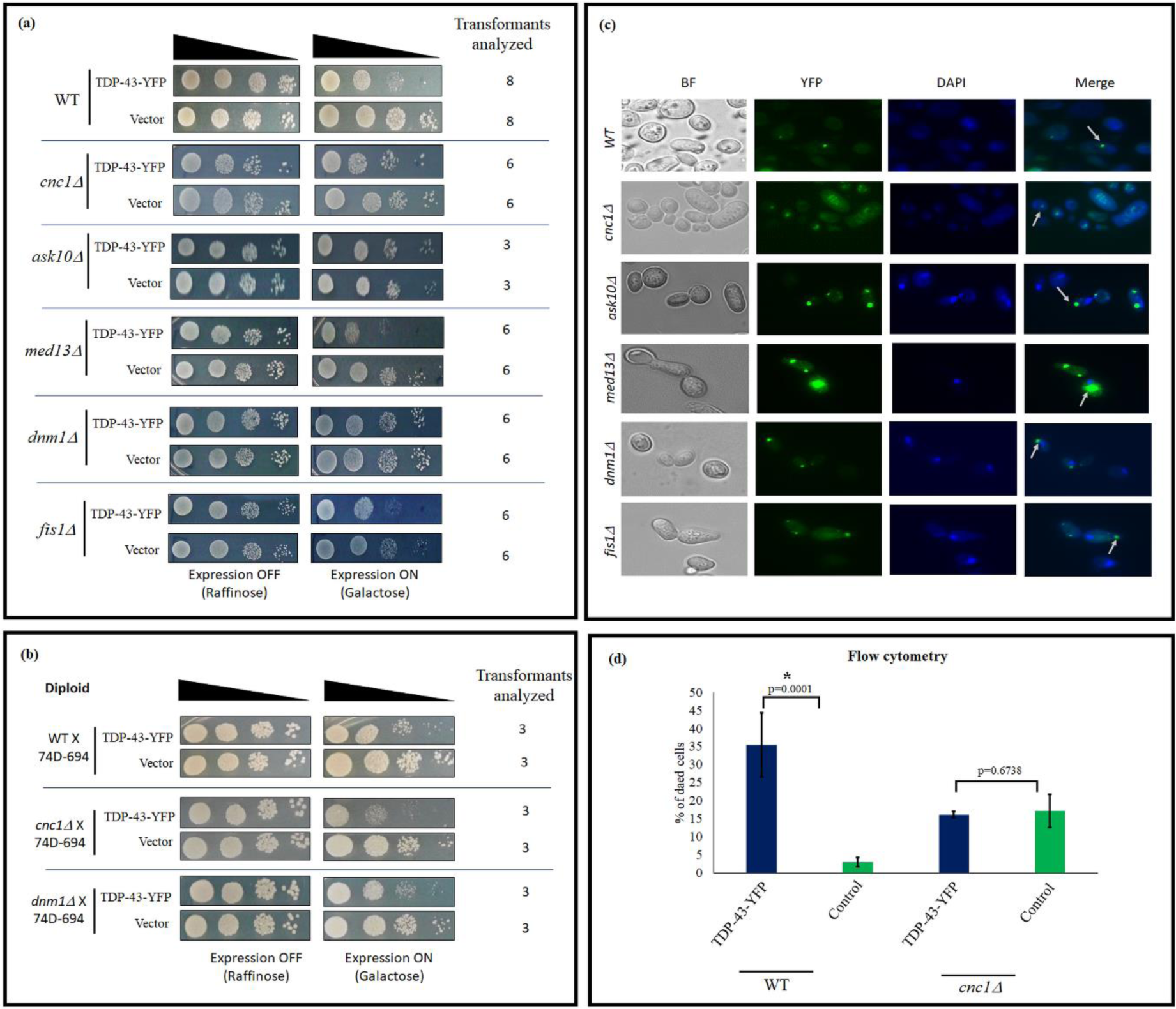
TDP-43 cytotoxicity in the yeast cells with deletion of the *CNC1* gene or its associated mitochondrial fission pathway genes. **a**. Wild-type (WT) yeast (BY4742) was transformed with low copy number, *GAL1* promoter-driven *CEN* plasmid encoding TDP-43-YFP (*pRS416-GAL1p-TDP-43-YFP, URA3*) or with the empty vector (*pRS416*). Serial dilution (10-fold) growth assays were performed onto plasmid-selective galactose-containing media where the TDP-43-YFP expression is turned on and also, for the cell number controls, onto raffinose-containing plates where the *GAL1* promoter is switched-off. Similar serial dilution (10-fold) growth assays were also performed for TDP-43-YFP plasmid transformants in the BY4742 derivative strains singly deleted for the gene, *CNC1*, encoding the Cyclin C protein or the proteins that function in conjunction with Cyclin C. The yeast strains deleted for *cnc1Δ, ask10Δ* or *dnm1Δ* showed significant rescue of the TDP-43-YFP toxicity. Whereas the yeast deleted for the genes *MED13* or *FIS1* manifested toxicity similar to that of the wild-type strain. At least three independent transformants in each deletion strain were analyzed to ensure consistency. The number of independent transformants examined for a particular deletion strain are indicated. **b.** For complementation assay, diploid strains were generated by crossing the haploid *cnc1Δ* or *dnm1Δ* or the wild-type (WT) in BY4742 background with a haploid WT strain (74D-694) of the opposite mating type. In the serial dilution (10-fold) growth assays, the obtained diploid strains pre-transformed and expressing the TDP-43-YFP, exhibited cytotoxicity thereby suggesting that the previously observed rescue of the toxicity was indeed due to the *bona fide* absence of the *CNC1* and *DNM1* genes in the BY4742 deletion parent strains. **c**. To examine if any of the gene deletions caused inhibition of the TDP-43-YFP aggregation or prevention of the cytoplasmic mis-localisation of TDP-43-YFP, the expression of TDP-43-YFP was induced with 2% galactose for 24 h in the gene deletion strains and the formation of the punctate fluorescent foci indicating of TDP-43-YFP aggregation, was examined under the YFP filter using Leica DM2500 microscope. To determine the sub-cellular location of the TDP-43-YFP aggregates, yeast cells were also stained with DAPI to visualize the nuclei and the localization of the aggregates with respect to the nucleus in a cell was examined. The presence of cytoplasmic TDP-43-YFP aggregates in each deletion strain has been marked using a white arrow. **d.** In order to corroborate the rescue of the TDP-43-YFP toxicity observed in the serial dilution assays, number of dead cells were counted using flow cytometry. For this, the yeast cells expressing TDP-43-YFP or control vector (EGFP) in the WT or *cnc1Δ* yeast, after 36 h of expression, were treated with propidium iodide (PI) and the quantitative measurement of the dead cells stained by PI were analysed by flow cytometry using FACS Aria III BD Bioscience machine. The data represent the mean number of dead cells from five independent transformants and the error bars represent standard deviation. The p-values were determined using un-paired t-test by comparing the vector controls (EGFP) with TDP-43-YFP in the WT or *cnc1Δ* yeast strain. The * in the p-value represents statistical significance.

Furthermore, to confirm that the effects observed on the TDP-43-YFP toxicity were indeed due to these gene deletions, and not due to any non-Mendelian genetic elements such as yeast prions that are known to affect heterologous amyloid protein aggregation and toxicity in yeast [61, 63], we further examined for complementation. For this, we first crossed the haploid wild-type *dnm1Δ* and *cnc1Δ* yeast with a wild-type haploid yeast strain of the opposing mating type and to obtain diploid strains and then expressed TDP-43-YFP in these diploids. Notably, the toxicity of TDP-43-YFP reappeared in all the diploid strains, thereby corroborating that the previously observed rescue of the toxicity was indeed due to the *bona fide* effects of the respective gene deletions (**Figure 2b**).

When there is low level of TDP-43 expression in the yeast, TDP-43 which harbours a nuclear localization signal (NLS) is known to be retained in the nucleus [45]. However, moderate/high expression of TDP-43 results in the formation of cytoplasmic aggregates recapitulating the TDP-43 pathology of cytoplasmic mis-localization of TDP-43 as also observed in the ALS patients. As the TDP-43-YFP toxicity was rescued in the *cnc1Δ, ask10Δ* and *dnm1Δ* yeast, we therefore examined whether there is any prevention of the aggregation of TDP-43 or its cytoplasmic mis-localization that maybe reducing the toxicity. However, we found that TDP-43-YFP formed cytoplasmic aggregates in all the deletion backgrounds similar to that of the wild-type (**Figure 2c**). Thus, the deletions of *CNC1, ASK10* and *DNM1* genes do not seem to modify the TDP-43 toxicity *via* preventing its cytoplasmic aggregation and possibly the rescue of toxicity is a consequence of defects in the downstream relay of the toxicity of the TDP-43 aggregates.

To confirm the results from the yeast growth assays and also to quantify the level of rescue, flow cytometry was employed to estimate cell death. When the TDP-43-YFP or the control vector-expressing cells were treated with propidium iodide (PI) and the dead cell population was counted using flow cytometry, TDP-43 expression in the wild-type caused significantly higher levels of cell death. On the contrary, and in conformity with the yeast growth assay, TDP-43 expression in the *cnc1Δ* yeast did not show any significant increase in the dead cell population as compared to that of the control (**Figure 2d** and **Supplementary Figure 1S)**.

### Deletion of the *CAF4* gene but not the *VPS1* gene of the peroxisome fission pathway rescues the TDP-43 toxicity

Notably, there are several common key players between the peroxisome fission and the mitochondrial fission processes. Both of these fissions in the yeast are mediated by the peroxisomal/mitochondrial membrane anchored protein Fis1 and the cytoplasmic protein Dnm1 [52]. Two adaptor proteins, Caf4 and Mdv1, help in anchoring Dnm1 to the peroxisomal/mitochondrial surfaces for effecting fission [55]. Independent of the Dnm1 pathway, the Vps1 protein also regulates the peroxisomal fission in yeast [55]. Both mitochondria and peroxisomes are known centres of the ROS generation and in the event of mitochondrial or peroxisomal hyper-fission the cellular longevity in yeast is compromised [53, 56]. Since, we observed the TDP-43-YFP toxicity was rescued in *dnm1Δ*, we then examined the TDP-43-YFP toxicity levels in the *caf4Δ, mdv1Δ* and *vps1Δ* yeast strains using growth assays. We found that there was a moderate rescue of the TDP-43-YFP toxicity in the *caf4Δ* and *mdv1Δ* whereas the *vps1Δ* strains exhibited the TDP-43-YFP toxicity at levels similar to that of the wild-type yeast (**Figure 3a**). Next, we examined whether there is any change in the aggregation pattern of TDP-43-YFP and found that TDP-43-YFP formed cytoplasmic aggregates in these deletion backgrounds similar to that of the wild-type (**Figure 3b**). Taken together, we observed rescue of the TDP-43 toxicity in the yeast deletions of the *DNM1, CAF4* and *MDV1* genes encoding for the proteins common to the mitochondrial and peroxisome fission pathways. On the contrary, the deletion of the *VPS1* gene which encodes for the protein, that is exclusively involved in the peroxisomal fission, did not affect the TDP-43 cytotoxicity.

**Figure 3:**
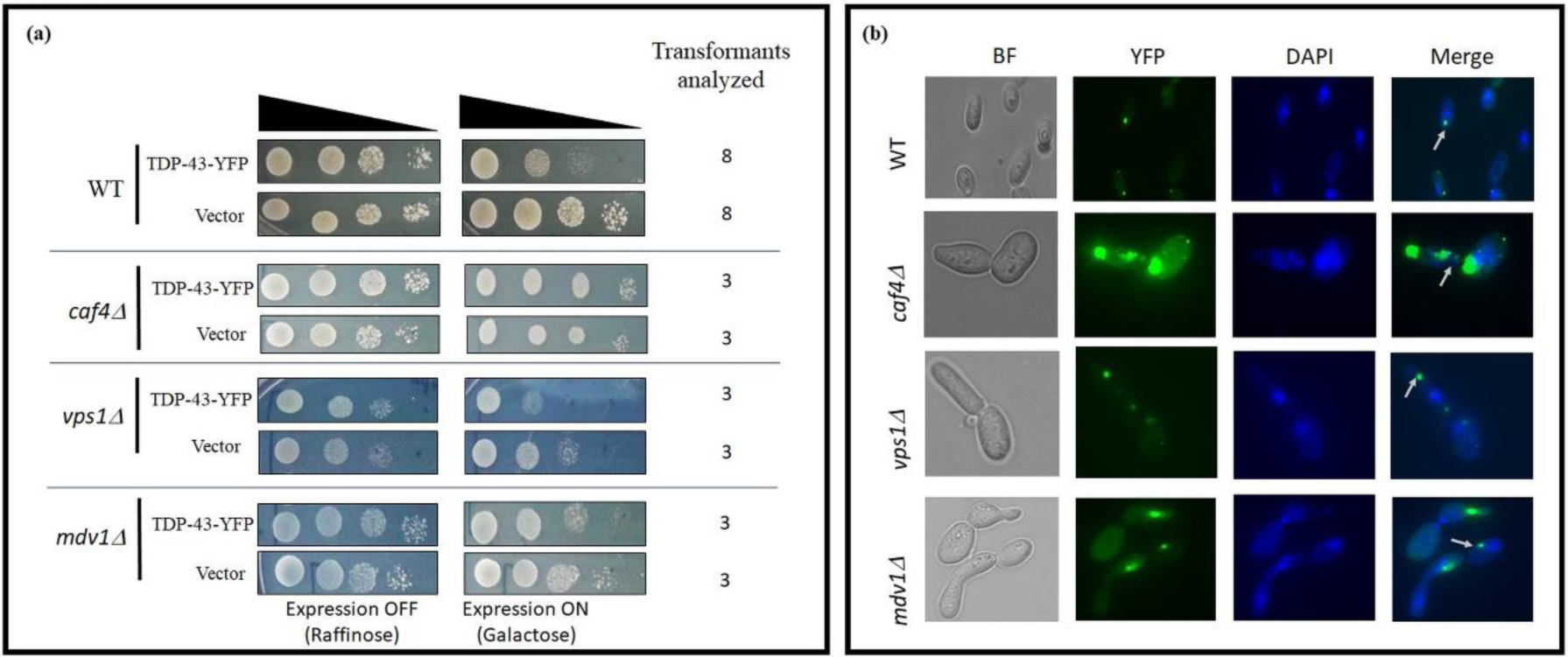
TDP-43 cytotoxicity in the yeast cells with deletion of the *CAF4, VPS1* and *MDV1* genes of the peroxisome fission pathway. **a**. Serial dilution (10-fold) growth assays were performed using the yeast strains deleted for the genes *CAF4, VPS1* or *MDV1* that code for proteins involved in the peroxisome fission pathway to examined cytotoxicity upon expression of TDP-43-YFP as compared to control vector (*pRS416*) transformants. The *caf4Δ* yeast strain showed significant rescue of the TDP-43-YFP toxicity and the *mdv1Δ* yeast manifested partial rescue, whereas, the *vps1Δ* yeast failed to rescue the TDP-43-YFP-induced cytotoxicity. At least three independent transformants were assessed to ensure that the observations are consistent. **b.** To ensure that the gene deletions do not interfere with the TDP-43-YFP aggregation or its cytoplasmic mis-localization, fluorescence microscopy was employed. TDP-43-YFP was expressed with 2% galactose for 24 h in the indicated gene deletion strains and the formation of the punctate TDP-43-YFP fluorescent foci was observed under YFP filter using Leica DM2500 microscope. To determine the sub-cellular location of the TDP-43-YFP aggregates, the yeast cells were also stained with DAPI to visualize the nuclei. Presence of cytoplasmic TDP-43-YFP aggregates are indicated with the white arrow in each gene deletion yeast strains.

### Deletion of the *CNC1* gene can even rescue the toxicities of highly overexpressed full-length TDP-43 and a C-terminal fragment TDP-43-(175-414)

As the Dnm1-independent peroxisomal fission modulator Vps1, did not appear to mediate the TDP-43 toxicity, we further examined how the TDP-43 toxicity is affected by *dnm1Δ* and *cnc1Δ* which are involved in the mitochondrial fission. As the toxicity resulted upon the low level expression from TDP-43-YFP encoded from a low copy *CEN* plasmid was rescued in the *dnm1Δ* and *cnc1Δ* yeast strains, thus we asked whether the toxicity of TDP-43 expressed at high levels from a multi-copy 2μ plasmid, can also be rescued in these deletion backgrounds. Thus, we first expressed TDP-43-GFP encoded from a 2μ plasmid driven by *GAL1* promoter and confirmed using western blotting that the TDP-43-GFP had several fold enhanced expression compared to TDP-43-YFP that was expressed from the low copy number plasmid (**Figure 4a** and **Supplementary Figure 2S**). As reported by the previous groups, the TDP-43-GFP over-expression exhibited enhanced toxicity in comparison with the TDP-43-YFP over-expression in the wild-type yeast strain. Notably, in the *dnm1Δ* yeast, we observed only a partial rescue of the TDP-43-GFP toxicity upon expression from the high copy number plasmid (**Figure 4b**). Expectedly, when there is enhanced expression and higher aggregation of TDP-43, some Dnm1-independent pathways may get involved in rendering the TDP-43 toxic (**Figure 4b**). Strikingly, however, in the *cnc1Δ* yeast, the toxicity of the overexpressed TDP-43-GFP was observed to be completely rescued. In fact, TDP-43-GFP toxicity rescue levels were similar to that of the TDP-43-YFP toxicity rescue levels in the *cnc1Δ* yeast. In summary, Cyclin C appears to be highly important and key mediator of the TDP-43 toxicity and therefore *cnc1Δ* yeast could rescue the toxicity of TDP-43 when expressed at low as well as high levels.

**Figure 4:**
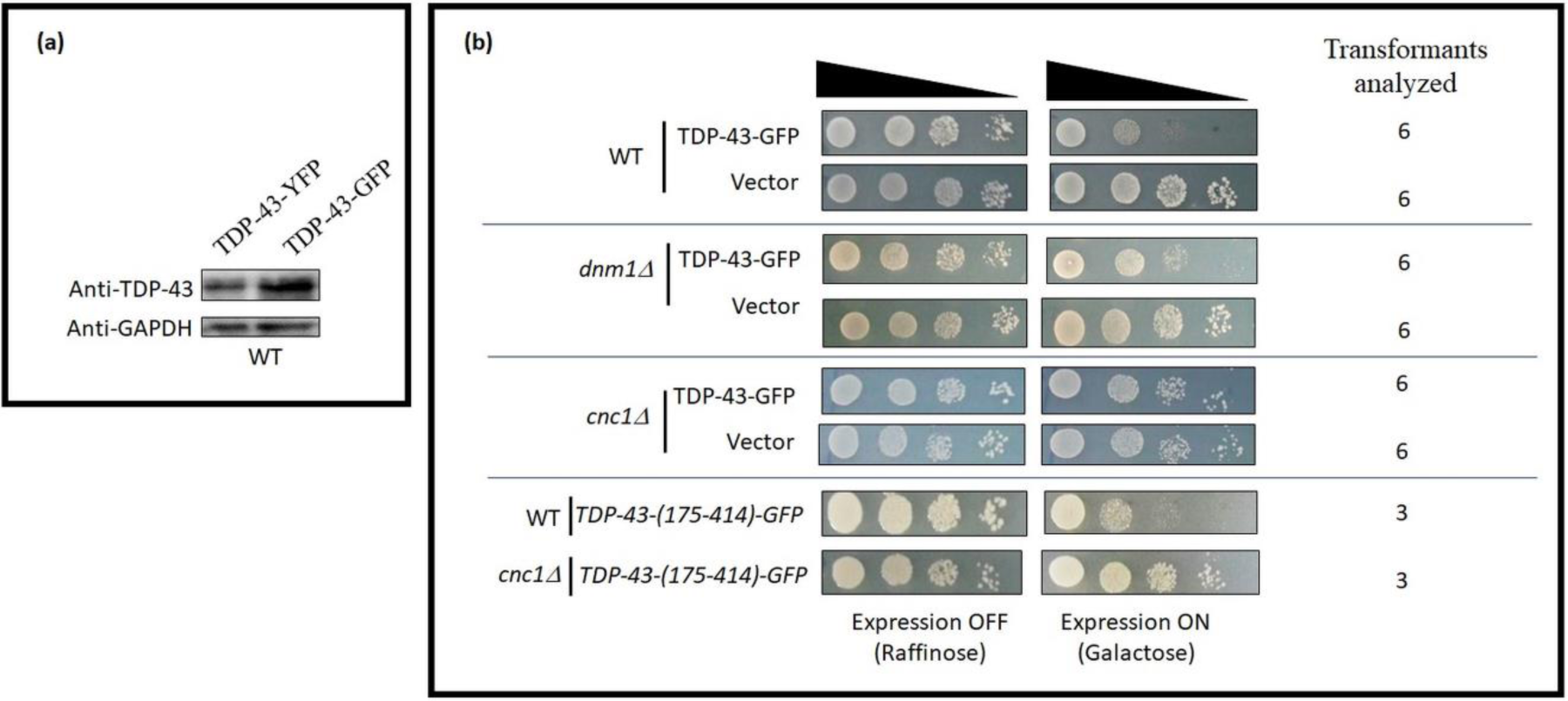
Deletion of the *CNC1* gene of the mitochondrial fission pathway can even rescue the toxicity of highly overexpressed full-length TDP-43 and a C-terminal fragment (aa:175-414) of TDP-43. **a**. Two fusion proteins, TDP-43-YFP expressed from a low copy number plasmid (*pRS416-TDP-43-YFP*) or TDP-43-GFP expressed from a high copy number plasmid (*pRS426-TDP-43-GFP*), both under *GAL1* promoter control, were first checked for relative TDP-43 protein expression levels after 24 h of expression by western blotting using anti-TDP-43 antibody. Endogenously expressed GAPDH protein was simultaneously probed with anti-GAPDH antibody, as the protein loading control. TDP-43-GFP exhibited many-fold higher expression than TDP-43-YFP. **b**. To examine if toxicity upon high level expression of TDP-43 can also be rescued, serial dilution (10-fold) growth assays were performed in the wild-type, *dnm1Δ* or *cnc1Δ* yeast strains expressing either the vector (*pRS416*) or TDP-43-GFP. In the *dnm1Δ* and *cnc1Δ* yeast there was a significant rescue of the TDP-43-GFP toxicity. Serial dilution (10-fold) growth assays were also performed in the wild-type and *cnc1Δ* strains expressing a toxic C-terminal fragment (CTF), usually generated by aberrant caspase cleavage of TDP-43 in the ALS patients, which we generated with GFP fusion (TDP-43-(175-414)-GFP) also in a high copy number plasmid under the *GAL1* promoter control. In the *cnc1Δ* yeast, there was a significant reduction in the toxicity of this CTF in comparison with that of the wild-type yeast strain. At least three independent plasmid transformants were analyzed in each case to confirm the observations.

Notably, several C-terminal fragments (CTFs) of TDP-43 have been previously found generated, proposedly by aberrant action of caspases, and deposited in the ALS patients. Also, these CTFs have also been proposed to be highly aggregation-prone and also projected to even have the capability of seeding the aggregation of the full-length TDP-43 [78, 79]. To examine whether the *cnc1Δ* yeast can rescue the toxicity of only the full-length TDP-43 or even that of the caspase generated CTF of TDP-43, we first generated a high copy plasmid encoding one such caspase generated CTF with aa: 175-414, which is generally referred to as the TDP-25 CTF [45, 79]. This fragment also encompasses the TDP-43 domains reported to be essential for the toxicity in the yeast [46]. We observed that the *cnc1Δ* yeast could rescue the toxicity of the overexpressed TDP-43-(175-414)-GFP similar to as observed previously for the full-length TDP-43-GFP (**Figure 4b**). This further supports of an important role of Cyclin C in mediating the TDP-43 toxicity and also indicates of similar mechanisms of the ensuing toxicities of the full-length TDP-43 and the TDP-25 CTF in the yeast cells.

### The *cnc1Δ* and *dnm1Δ* yeast strains can rescue the TDP-43 toxicity only in the cells with functional mitochondria

Earlier, the presence of the functional mitochondria has been shown to be important for mediating the TDP-43 toxicity in yeast [22]. Recently, respiration was found to enhance the TDP-43 toxicity, but the toxicity was not found to be completely abrogated in the absence of respiration [80]. The Cyclin C protein functions in both transcriptional regulation as well as mitochondria-mediated cell death pathway. Thus, to dissect out whether the observed rescue of the TDP-43 toxicity in the *cnc1Δ* yeast, is due to the loss of the Cyclin C’s function related to the transcriptional regulation or due to the loss of the mitochondria-related role, we generated petite strain, [*rho*^*-*^], of the *cnc1Δ* yeast and also for comparison that of the wild-type and *dnm1Δ* yeast strains. The [*rho*^*-*^] strains have non-functional mitochondria which are compromised for respiratory capability. Then, we examined the rescue of TDP-43 toxicity in the *cnc1Δ* and *dnm1Δ* yeast with either functional [*RHO*^*+*^] or non-functional mitochondria [*rho*^*-*^] (**Supplementary Figure 3S**). For this, we used the *GAL1* promoter-driven low copy number plasmid encoded expression of TDP-43-YFP in the *cnc1Δ, dnm1Δ* and wild-type yeast strains, which are all derivatives of the BY4742 strain. While several yeast strains such as 74-D-694 when compromised for the mitochondrial respiration do not utilize galactose, the respiration-deficient derivatives of BY4742 are known to successfully utilize galactose [81, 82]. Notably, in contrast to the observed significant reduction in the TDP-43-YFP toxicity in the *cnc1Δ* and *dnm1Δ* yeast cells containing the respiring and functional mitochondria (i.e. [*RHO*^*+*^] yeast), in the isogenic yeast strains with non-respiring mitochondria (i.e. [*rho*^*-*^] yeast), the TDP-43-YFP expression-caused toxicity was not rescued (**Figure 5**). Possibly, toxicity mechanisms independent of the mitochondrial fragmentation dominate and mediate the TDP-43 toxicity in the absence of the functional mitochondria. This lack of rescue of the TDP-43 toxicity in the [*rho*^*-*^] version of the *cnc1Δ* yeast rules out the involvement of transcriptional role of Cyclin C in mediating the TDP-43 toxicity. Rather, it implicates the non-canonical non-transcriptional role of Cyclin C, which is mediated through the functional mitochondria, in affecting the toxicity of TDP-43 in the yeast cells.

**Figure 5:**
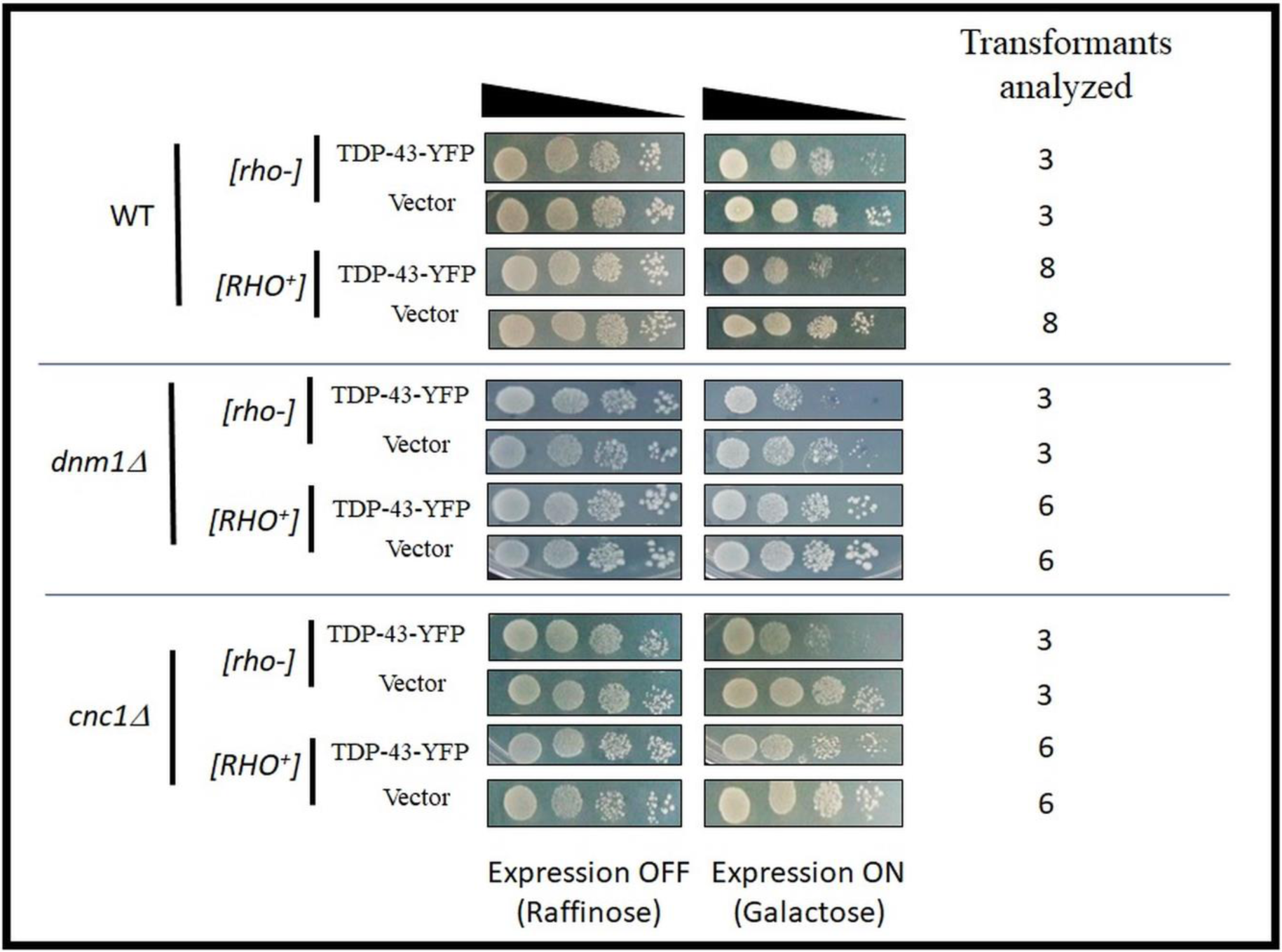
TDP-43-YFP cytotoxicity in the *dnm1Δ* and *cnc1Δ* yeast strains containing functional or non-functional mitochondria. To check whether the effects of *CNC1* gene on the TDP-43 toxicity was mediated *via* the mitochondria-dependent pathway. Isogeneic non-respiring [*rho*^*-*^] versions of the wild-type (WT), *dnm1Δ* and *cnc1Δ* strains were pre-generated by growing the strains on YPD with 0.4 mg/mL of ethidium bromide and then transformed with TDP-43-YFP encoding plasmid (*pRS416-GAL1p-TDP-43-YFP*). Serial dilution (10-fold) growth assays were performed in the isogenic WT, *dnm1Δ* and *cnc1Δ* yeast strains expressing either the vector (*pRS416*) or TDP-43-YFP in the yeast strains with functional mitochondria [*RHO*^*+*^] or non-functional mitochondria [*rho*^*-*^]. At least three independent plasmid transformants in each case were investigated simultaneously to ensure the consistency in the observations.

### Deletion of the yeast pro-apoptotic gene *YBH3* rescues the TDP-43 toxicity

Previous study has shown that the TDP-43-expressing yeast cells display markedly increased apoptotic markers, however the key mitochondrial cell death proteins such as apoptosis inducing factor or cytochrome c were not to found to directly mediate the cell death upon the TDP-43 expression [22]. Therefore, considerable interest exists in finding out the events and key players that culminate the mitochondria-dependent cell death upon TDP-43 expression in yeast. It is known that the permeabilization of the mitochondrial outer membrane is mediated by BH3 domain-containing proteins and the yeast BH3 containing protein, Ybh3, is involved in mediating mitochondria-dependent apoptosis and the deletion of the *YBH3* gene confers resistance to the apoptosis induction [22]. As our results showed that the deletions of certain genes involved in the mitochondrial fission pathway rescue the TDP-43 toxicity, we examined whether *ybh3Δ* yeast, where there is a loss of the pro-apoptotic protein Ybh3, has any effect on the TDP-43 toxicity. The Ybh3 protein, which is generally a vacuolar protein, translocates to the mitochondria upon oxidative stress and induces the programmed cell death. Interestingly, we found that the deletion of the *YBH3* gene can alleviate the TDP-43-YFP-induced cytotoxicity and cell death (**Figure 6**). In addition, the toxicity of the highly overexpressed TDP-43-GFP was also partially rescued in the *ybh3Δ* yeast (**Supplementary Figure 4S**). Furthermore, in coherence with the previous results on the *cnc1Δ* yeast, we also found that even though there was a significant reduction in the TDP-43-YFP toxicity in the *ybh3Δ* yeast cells containing the respiring and functional mitochondria (i.e. [*RHO*^*+*^] yeast), in the isogenic yeast strains with non-respiring mitochondria (i.e. [*rho*^*-*^] yeast), the TDP-43-YFP expression continued to manifest toxicity (**Figure 6**). Next, to confirm that the observed rescue of the TDP-43-YFP toxicity was indeed due to the lack of Ybh3 function in the *ybh3Δ* yeast and not due to any non-Mendelian genetic elements, we examined for complementation. For this, we crossed the haploid *ybh3Δ* yeast, with a wild-type haploid 74-D-694 yeast strain of the opposing mating type to obtain diploids and then transformed and expressed TDP-43-YFP. Notably, the toxicity of TDP-43-YFP reappeared in the diploid strain, thereby corroborating that the previously observed rescue of the toxicity was indeed due to the *bona fide* absence of the Ybh3 function in the *ybh3Δ* yeast (**Figure 6b**).

**Figure 6:**
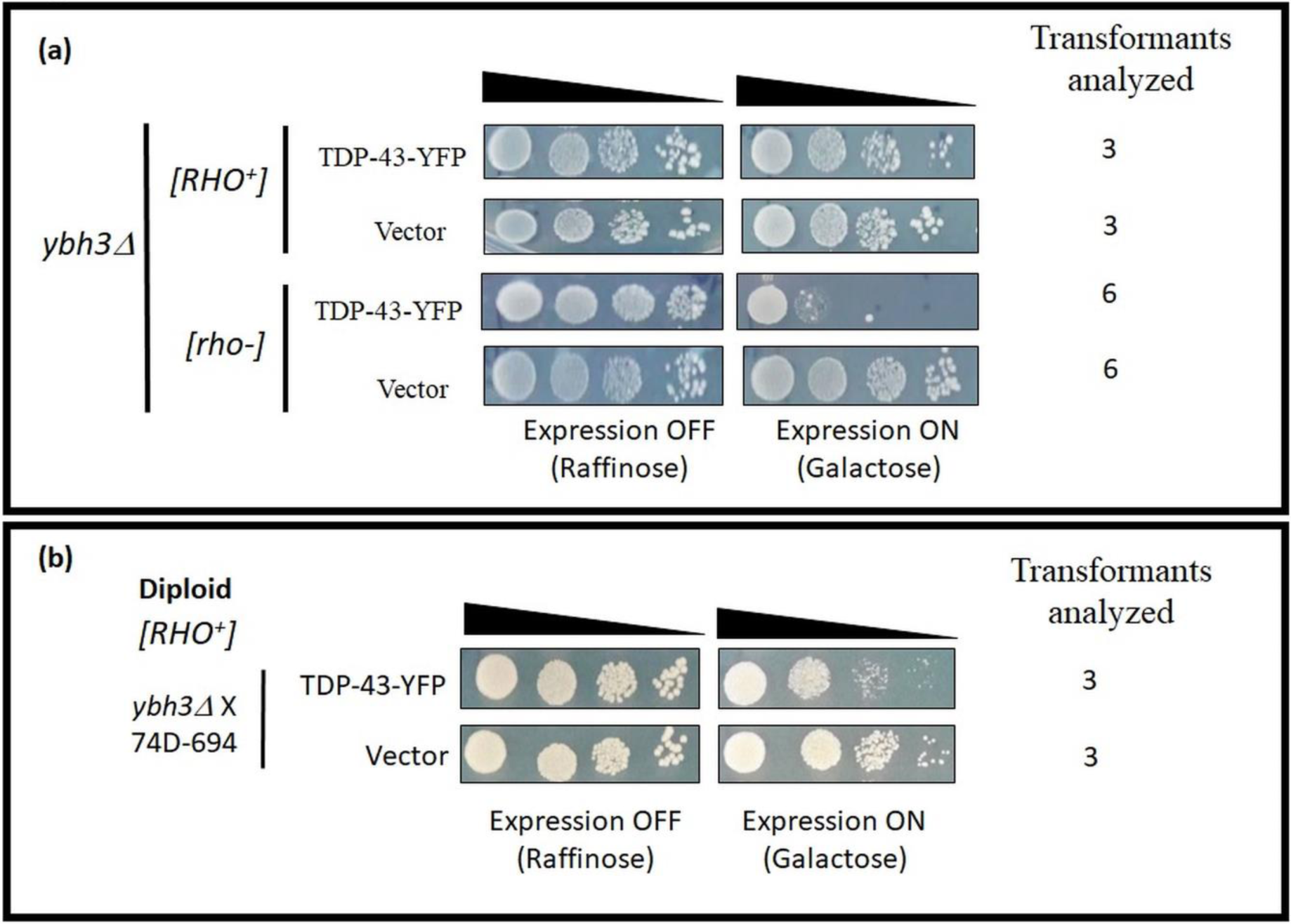
TDP-43 cytotoxicity in the yeast cells with deletion of the pro-apoptotic gene *YBH3*. **a**. To examine whether the cytotoxicity of TDP-43 depends on the presence of the pro-apoptotic gene *YBH3* and whether functional mitochondria is required for it effect, we first obtained a petite version of [*rho*^*-*^] of the BY4742 derivative *ybh3Δ* yeast strain and then transformed it with TDP-43-YFP encoding plasmid (*pRS416-GAL1p-TDP-43-YFP*). Serial dilution (10-fold) growth assays of the [*RHO*^*+*^] or [*rho*^*-*^] transformants in the *ybh3Δ* yeast, were performed to examine the TDP-43-YFP toxicity compared to the vector control (*pRS416*). The yeast strain *ybh3Δ* showed significant rescue of the TDP-43-YFP toxicity only when functional mitochondria were present. **b.** For complementation assay, diploid strains were generated by crossing the [*RHO*^*+*^] haploid *ybh3Δ* yeast in the BY4742 background with a haploid wild-type strain (74D-694) of the opposite mating type. In serial dilution (10-fold) growth assays, the diploid strain expressing the TDP-43-YFP, exhibited toxicity thereby suggesting that the previously observed rescue of toxicity was indeed due to the *bona fide* absence of the *YBH3* gene in the deletion parent strains. At least three independent plasmid transformants in each case were investigated simultaneously to corroborate the consistency in the observations.

### Although TDP-43 aggregation-induced ROS is present in the *cnc1Δ* and *dnm1Δ* yeast cells, the cell death is prevented

As the expression of TDP-43 is known to elicit oxidative stress in the wild-type yeast [22, 47, 48], we examined here the levels of reactive oxygen species (ROS) in the *cnc1Δ* and *dnm1Δ* yeast using the cell permeant reporter dye CellROX deep red. This dye usually remains non-fluorescent in the absence of ROS but exhibits bright red fluorescence upon oxidation by ROS. In the wild-type cells, upon TDP-43-YFP expression for 36 h, about 16% of the cells exhibited bright red fluorescence whereas only ∼1% of the cells exhibited CellROX staining in the vector controls. This corroborates the previously documented high ROS generation upon TDP-43 expression in yeast. Next, we examined whether TDP-43-YFP expression also induces ROS in the yeast deleted for the mitochondrial fission genes, *CNC1* and *DNM1*, or the observed rescue of the TDP-43-YFP toxicity in these backgrounds is due to the absence of ROS generation. When counted using flow cytometry, we found that only around 7% of the TDP-43-YFP-expressing cells and around 1% the of vector transformants exhibited the CellROX red fluorescence in the *dnm1Δ* yeast thereby suggesting of an overall reduction in the ROS levels relative to the wild-type yeast but however, not a complete abrogation of the ROS generation **(Figure 7)**. Also, the ROS-positive cell numbers were statistically significantly higher in the TDP-43-YFP-expressing samples in comparison with those in the vector transformed *dnm1Δ*. Notably, around 17% of cells among the *cnc1Δ* yeast cells with TDP-43-YFP expression and ∼1% of the cells with the vector exhibited the fluorescence of CellROX, thereby indicating a lack of any overall reduction in the ROS levels in the *cnc1Δ* yeast compared to the wild-type yeast cells (**Figure 7**). This data suggests that although TDP-43 expression induces considerable ROS levels in the *cnc1Δ* and *dnm1Δ* yeast, the lack of the Dnm1 and Cyclin C proteins prevents the ROS from culminating into the cell death.

**Figure 7:**
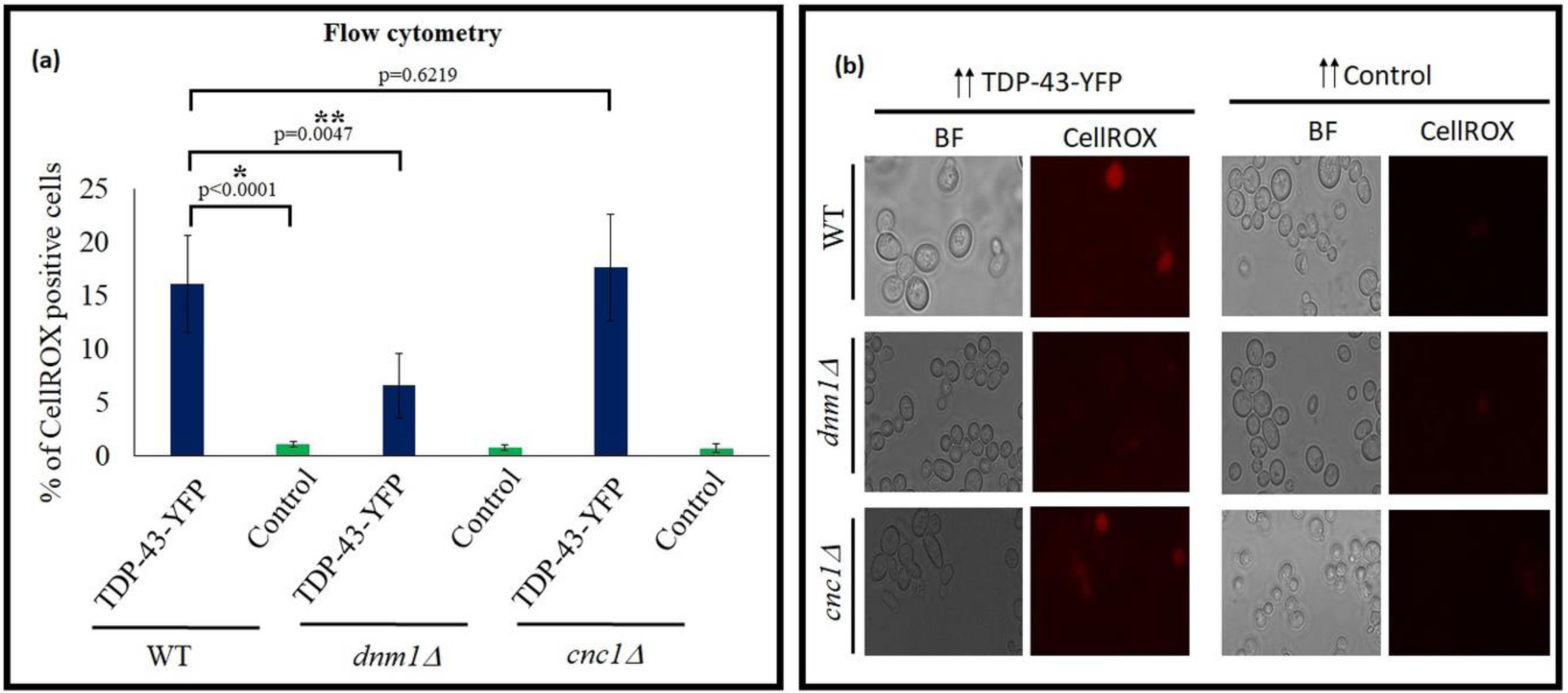
Assessment of the ROS levels in the yeast cells deleted for the *DNM1* and *CNC1* genes. To test whether the *cnc1Δ* and *dnm1Δ* yeast that show the rescue of TDP-43 toxicity fail to accumulate ROS upon TDP-43 expression or rather fail to transmit the effect of ROS into cell death, any presence of ROS in the yeast cells expressing TDP-43-YFP was visualized by the reporter CellROX deep red reagent. CellROX is generally non-fluorescent but converted to bright red fluorescent species in the presence of ROS. The yeast cells were grown in 2% galactose for 24 h for expressing the TDP-43-YFP or the vector control (EGFP) before treatment with CellROX deep red reagent. **a.** The quantitative measurement by flow cytometry of the cells stained by CellROX. The data represent the mean number of CellROX-positive cells from five independent transformants and the error bars represent standard deviation. The p-values were determined with un-paired t-test by comparing the TDP-43-YFP in *cnc1Δ* or *dnm1Δ* using the TDP-43-YFP in the wild-type yeast. In the wild-type yeast, the expression of TDP-43-YFP showed significant increase in the number of cells manifesting CellROX fluorescence compared to the vector control (EGFP). However, the number of cells exhibiting fluorescence upon the TDP-43-FFP expression was markedly decreased in the yeast deleted for mitochondrial fission gene, *dnm1Δ.* On the contrary, the *cnc1Δ* yeast did not show difference in the oxidative stress levels as compared to that of the wild-type strain. The * on the p-value represents statistical significance between CellROX-positive cells of TDP-43-YFP expressing or the vector control cells in wild-type. ** represents statistical significance between CellROX-positive cells due to TDP-43-YFP expression in wild-type and *dnm1Δ*. **b.** Yeast cells expressing TDP-43-YFP or the vector control (EGFP) as described above in the wild-type, *cnc1Δ* or *dnm1Δ* were treated with CellROX and visualized under RFP filter using Leica DM2500 fluorescence microscope.

### TDP-43-induced oxidative stress causes the movement of Cyclin C from nucleus to the cytoplasm

Cyclin C is a predominantly a nuclear protein that translocates to the cytoplasm in the event of oxidative stress [57, 83]. Cytoplasmic localization of Cyclin C upon stress is observed from yeast to mammals and it results in the induction of the programmed cell death [84, 85]. Recently, it has been shown in the mammalian cell lines that the cytoplasmic Cyclin C protein directly stimulates the GTP affinity of the Drp1 protein and promotes mitochondrial fission in response to oxidative stress [84]. As the TDP-43 expression results in the enhanced oxidative stress and we observed that the TDP-43 toxicity is rescued in the *cnc1Δ* yeast, we therefore, examined whether there are any alterations in the sub-cellular localization of Cyclin C in yeast upon the TDP-43 expression. In a previous study, to examine the cellular localization of Cyclin C in proximity to the mitochondria upon oxidative stress, the Cyclin C protein tagged with YFP and a mitochondria localizing protein tagged with DsRed, were used suggesting a lack of interference between these two types of fluorescence [75]. On the similar lines, we constructed a Cyclin C-YFP expressing plasmid and used it in conjunction with a plasmid encoding TDP-43 DsRed. Similar to the observed rescue of the TDP-43-YFP and TDP-43-GFP toxicity in the *cnc1Δ* yeast, we first confirmed of a significant reduction in the TDP-43-DsRed toxicity in the *cnc1Δ* yeast (**Figure 8a**). In the control sample where the TDP-43-DsRed expression was absent, as expected Cyclin C-YFP was found to be localized mostly to the nucleus and only ∼3 % of the cells exhibited Cyclin C-YFP localized to the cytoplasm (**Figure 8b, 8c** and **Supplementary Figure 5S**). On the contrary, when TDP-43-DsRed was also co-expressed with Cyclin C-YFP, ∼30% of the cells exhibited cytoplasmic localization of Cyclin C-YFP. Furthermore, we also examined whether an untagged TDP-43 protein’s co-expression along with Cyclin C-YFP can also drive the cytoplasmic translocation of Cyclin C-YFP similar to the observations with the TDP-43-DsRed’s co-expression. For this we used 74-D-694 yeast strain and co-transformed it with *pGAL1-TDP-43 (TRP1)* and *pRS426-GAL1p-CNC1-YFP* (*URA3*). Indeed, significant percentage of the yeast cells were found to show Cyclin-C-YFP in the cytoplasm when it was co-expressed with the untagged TDP-43 as opposed to the vector co-expression controls (Data not shown). Recently, it has been shown that the TDP-43 toxicity in the yeast is significantly reduced in the presence of the reducing agent, N-acetyl cysteine (NAC) [80]. We first checked and confirmed that the TDP-43-induced oxidative stress was also significantly lowered when cells were grown with the addition of NAC (**Supplementary Figure 6S)**. Strikingly, when the cells co-expressing Cyclin C-YFP and TDP-43-DsRed were grown in presence of the anti-oxidant, NAC, the cytoplasmic localization of Cyclin C was significantly thwarted (**Figure 8b, 8c**, and **Supplementary Figure 6S**). This suggests that the TDP-43 expression-induced ROS-mediated translocation of Cyclin C from the nucleus to cytoplasm is an important mediator of the TDP-43-induced cytotoxicity and the consequent cell death in yeast.

**Figure 8:**
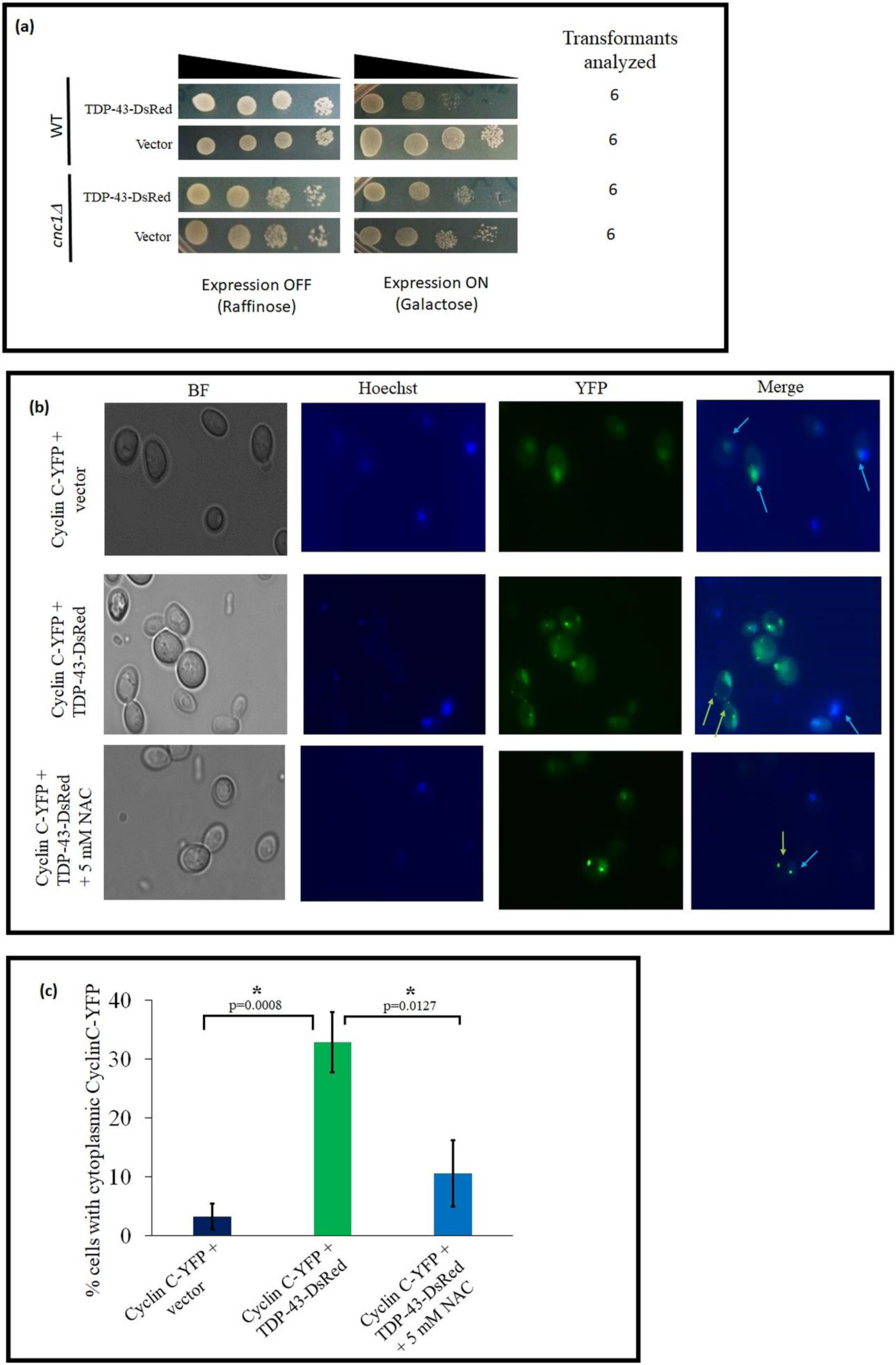
Oxidative stress-induced cytoplasmic localization of Cyclin C upon TDP-43 co-expression. **a**. Serial dilution (10-fold) growth assays of the wild-type and the yeast strain deleted for the gene *CNC1* upon the expression of TDP-43-DsRed from plasmid (*pGAL1-TDP-43-DsRed*) The *cnc1Δ* yeast strain showed significant rescue of the TDP-43-DsRed toxicity. **b** and **c.** Cyclin C-YFP and TDP-43-DsRed, both under *GAL1* promoter, were expressed with 2% galactose for 18 h in the wild-type BY4742 yeast. The cells were then stained with Hoechst stain for visualizing the nucleus. When examined by fluorescence microscopy, in the absence of TDP-43-DsRed expression, Cyclin C-YFP exhibited predominantly nuclear localization whereas upon TDP-43-DsRed co-expression, significant number of cells manifested presence of Cyclin C-YFP in the cytoplasm. Blue arrows indicate nuclear Cyclin C-YFP whereas yellow arrows point to cytoplasmic Cyclin C-YFP. When an anti-oxidant, N-acetyl-cysteine (NAC) was added in the cells co-expressing both Cyclin C-YFP and TDP-43-DsRed, the cytoplasmic translocation of Cyclin C-YFP was significantly reduced. Blindly acquired images from three independent transformants were manually counted for presence of cytoplasmic Cyclin C-YFP using only the yeast cells that exhibited both the nuclear staining by Hoechst and also the expression of Cyclin C-YFP. The error bars represent standard deviation. The p-values were determined with un-paired t-test. * in the p-value represents statistical significance.

## DISCUSSION

Previously, the TDP-43 overexpression in the yeast cell has been shown to mimic ALS-like cytoplasmic mis-localization of TDP-43 and cause cytotoxicity and furthermore, several yeast genes and metabolic pathways have been implicated in mediating the TDP-43 toxicity in the yeast cell [20, 46, 68, 69, 86]. Overall, several molecular mechanisms have been proposed for causing the TDP-43 proteinopathy such as impaired endocytosis, increased localization of TDP-43 to the mitochondria, dysregulation of the ubiquitin proteasome-mediated protein degradation, prion-like behaviour of TDP-43, disturbances in the chromatin remodelling and oxidative stress etc. [2, 11, 16-22, 87]. Of note, TDP-43 has also been shown to aberrantly localize to the mitochondria and affect the mitochondrial translation [19]. Additionally, it has been shown that the TDP-43 expression exacerbates the mitochondrial damage *via* the activation of the mitochondrial unfolding protein response pathway in both the cellular and animal models and proposedly, reversing or blocking the mitochondrial damage could potentially alleviate the TDP-43-mediated neurodegeneration [19]. In the single cell eukaryotic yeast model, the TDP-43 expression has been shown to induce oxidative stress and mitochondria have been implicated in mediating the TDP-43 toxicity, however the genes/proteins that are the key players, remain to be unravelled [22, 47, 48]. In fact, deletion of neither the mitochondrial apoptosis inducing factor gene nor the cytochrome c gene could relieve the TDP-43 toxicity although the yeast cells expressing TDP-43 displayed apoptotic markers [22]. Thus, here we examined further to learn how the cell death ensues upon the TDP-43 expression by investigating whether the oxidative stress induced by TDP-43 expression or the presence of mitochondrial fission genes or a pro-apoptotic gene, play any role.

Here, we find that the deletion of the *CNC1* gene, which is known to be involved in the oxidative stress response and the mitochondrial hyper fission, markedly decreased the TDP-43 toxicity. As the Cyclin C-mediated oxidative stress response is a conserved phenomenon from the yeast to humans, the observed rescue of the TDP-43 toxicity in the *cnc1Δ* yeast could be of high relevance to the ALS pathology. Cyclin C has been found to be associated with the Cdk8 protein in the nucleus and their interaction is essential for the transcriptional repression of the stress response genes in the unstressed cells. As we observed that the TDP-43 toxicity was rescued in the *cnc1Δ* yeast only when they contained functional mitochondria (i.e. [*RHO*^*+*^]) but not when they contained non-functional mitochondria (i.e. [*rho*^*-*^]), the transcriptional role of Cyclin C in mediating the TDP-43 toxicity is ruled out. Rather, the data support mitochondria-dependent Cyclin C’s role in mediating the TDP-43 toxicity possibly *via* mitochondrial hyper fission-mediated apoptosis, as the role of Cyclin C in mediating the mitochondrial hyper fission is established in the yeast cells [57, 83, 84].

Recently, it has been shown that Cyclin C, when present in the cytoplasm, directly stimulates the affinity of Drp1 (human homolog of Dnm1) for GTP for mediating the mitochondrial fission [84]. Here, we observed that the deletion of *DNM1* also markedly relieved the TDP-43 toxicity. Alike to the *cnc1Δ* yeast presence of functional mitochondria was also essential for the rescue in the *dnm1Δ* yeast. It is noteworthy that an inhibitor of Drp1 and Fis1 interaction was previously found to relieve the TDP-43-induced mitochondrial fragmentation and cytotoxicity [88]. Surprisingly however, the toxicity of TDP-43 was not rescued in the *fis1Δ* yeast the reason for which is not completely understood. Notably, it is also previously documented that *fis1Δ* yeast is still capable of undergoing mitochondrial fragmentation which may be a reason for the lack of the TDP-43 toxicity rescue in the *fis1Δ* yeast [56]. We also observed here that the deletion of the *MED13* gene that encodes for the Med13 protein which is essential for the nuclear retention of Cyclin C failed to rescue the TDP-43 toxicity [89, 90]. On the contrary, the deletion of the *ASK10* gene which codes for the Ask10 protein that is responsible for the cytoplasmic re-localization of Cyclin C, was found here to significantly reduce the TDP-43 toxicity. Taken together, the factors influencing the cytoplasmic translocation of Cyclin C seem vital in mediating the TDP-43 toxicity.

As there is a considerable overlap in the array of proteins involved in the mitochondrial and peroxisomal fission processes, we also examined the TDP-43 toxicity in the yeast strains deleted for the genes encoding proteins involved in the peroxisomal pathway. We find that the toxicity of TDP-43 is substantially rescued in the *dnm1Δ* and *caf4Δ* and partially recued in the *mdv1Δ* yeast strains. All of these deletion strains lack a protein that contributes both to the fission of the mitochondria as well as that of the peroxisomes. On the contrary, the deletion of *VPS1* gene that encodes for the Vps1 protein which is an independent regulator of the peroxisomal fission failed to rescue the TDP-43 toxicity. Thus, peroxisome fission seems an unlikely key player in the TDP-43-mediated cytotoxicity and cell death in yeast.

Remarkably, the *cnc1Δ* yeast could rescue the toxicity of TDP-43 when it was expressed at low or even high levels as well as irrespective of whether the TDP-43 protein was fused to YFP, GFP or DsRed. Even the toxicity of a CTF, TDP-25, was also rescued in the *cnc1Δ* yeast. Thus, the data support of Cyclin C being a *bona fide* key player in mediating the toxicity of TDP-43 in the yeast cells.

It is known that the mitochondrial localization of the yeast pro-apoptotic protein, Ybh3, which contains Bcl-2 homology domain (BH3) similar to that of the several human pro-apoptotic proteins, is needed to initiate the mitochondria-mediated programmed cell death (PCD) in yeast [22, 57]. Strikingly, we found here that the loss of the *YBH3* gene that encodes for the Ybh3 protein also rescues the TDP-43-induced cytotoxicity. As this rescue was observed only in the [*RHO*^*+*^] yeast but not in the [*rho*^*-*^] yeast, therefore the mitochondria-dependent, Ybh3-mediated PCD appears important for the TDP-43-induced cytotoxicity in yeast.

Furthermore, although the TDP-43-induced oxidative stress was present in the wild-type, *cnc1Δ* and *dnm1Δ* yeast strains, but the cell death was significantly reduced in the *cnc1Δ* and *dnm1Δ* yeast thereby suggesting of a missing relay link between the ROS and the downstream PCD. If Cyclin C was the link to relay the TDP-43-induced ROS effect to culminate into cell death, it would be expected that Cyclin C moves to the cytoplasm upon the TDP-43 expression. Indeed, Cyclin C-YFP was observed to translocate from the nucleus to the cytoplasm upon the TDP-43 co-expression. This strongly suggests that the oxidative stress caused by the TDP-43 expression could relay the toxicity and cause cell death *via* Cyclin C. In accordance, the addition of the anti-oxidant molecule N-acetyl-cysteine (NAC), which has been previously shown to relieve the TDP-43 toxicity in the yeast [80], also significantly prevented the cytoplasmic localization of Cyclin C-YFP. Recently, it has been shown that Cyclin C associates with Bcl-2 homology containing protein BAX and tethers it with the stress-induced mitochondrial fission complex in the mammalian cell line. This provides sufficient dwell time on mitochondria for BAX to be recognized by the BH3 containing proteins leading to initiation of PCD in the mammalian cell line [88]. Since Cyclin C translocation from the nucleus to the cytoplasm is a conserved process from the yeast to humans, this finding of TDP-43-induced Cyclin C translocation in yeast may have direct relevance to the ALS pathology [83, 85, 88, 91, 92]. Therefore, targeting the TDP-43 expression-induced oxidative stress and the Cyclin C translocation may have potentially fruitful consequences towards finding of the ALS therapeutic targets.

## ACKNOWLEDGEMENTS

We sincerely thank Prof. Susan W Liebman, University of Nevada Reno, USA, for kindly gifting yeast strains and plasmids. We also thank Dr. Parag Pawar and Mr. Ch. V. Tejesh Reddy, IIT Hyderabad for help with flow cytometry. We thank IIT Hyderabad funded by MHRD, Govt. of India for research infrastructure and support. VB thanks DBT, Govt. of India, for senior research fellowship (SRF). AG is thankful to MHRD, Govt. of India, for SRF. Basant K Patel thanks DST, Govt. of India for research grant (No: EMR/2016/006327).

