## Supplementary Figures for "Role of *CNC1* gene in TDP-43 aggregation-induced oxidative stress-mediated cell death in *S. cerevisiae* model of ALS"

**Title**

**Supplementary Figure 1S**: **Cytotoxicity of TDP-43-YFP in wild-type and *cnc1Δ* yeast measured by flow cytometry**

*S. cerevisiae* wild-type or *cnc1Δ* cells expressing TDP43-YFP or the control (EGFP) were induced using 2% galactose for 36 hours. More than 1 million cells were harvested, washed and stained with propidium iodide and incubated for 15 minutes in the dark. 50,000 cells were acquired in AriaIII FACS (BD) using PE-Texas Red-A (for PI). Percentage of dead cells indicate that the TDP-43-YFP mediated toxicity is more in the wild-type yeast as compared to the *cnc1Δ* yeast.

#### Supplementary Figure 1S

Wild-type

TDP-43-YFP

Control

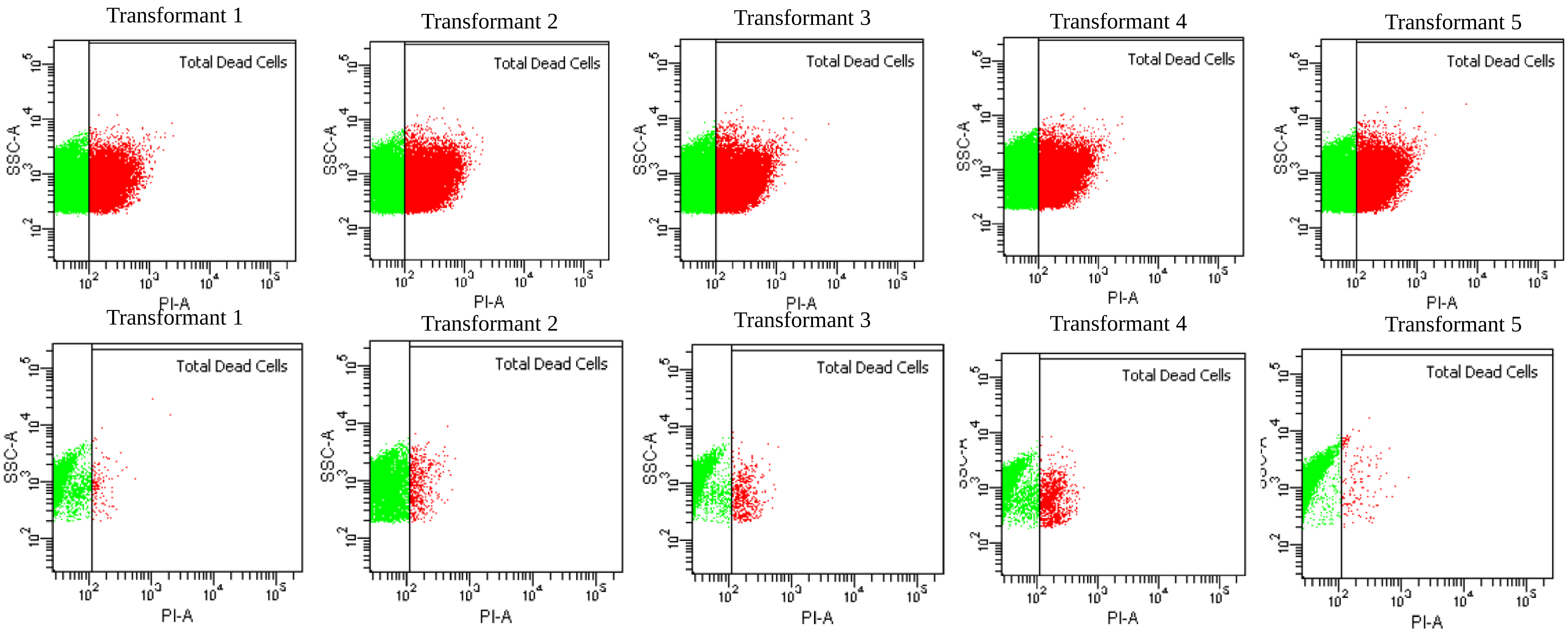

| | | Transformant 1 | Transformant 2 | Transformant 3 | Transformant 4 | Transformant 5 |
|---|---|---|---|---|---|---|
| | Total cells | 50000 | 50000 | 50000 | 50000 | 50000 |
| | Dead cells | 11251 | 20581 | 21109 | 15000 | 20734 |
| Wild-type TDP-43-YFP | % of Dead cells | 22.5 | 41.2 | 42.2 | 30 | 41.5 |
| | | Transformant 1 | Transformant 2 | Transformant 3 | Transformant 4 | Transformant 5 |
| | Total cells | 50000 | 50000 | 50000 | 50000 | 50000 |
| | Dead cells | 1570 | 2005 | 1485 | 2152 | 523 |
| Wild-type control | % of Dead cells | 3.14 | 4.01 | 2.97 | 4.304 | 1.046 |

#### Supplementary Figure 1S

*cnc1Δ*

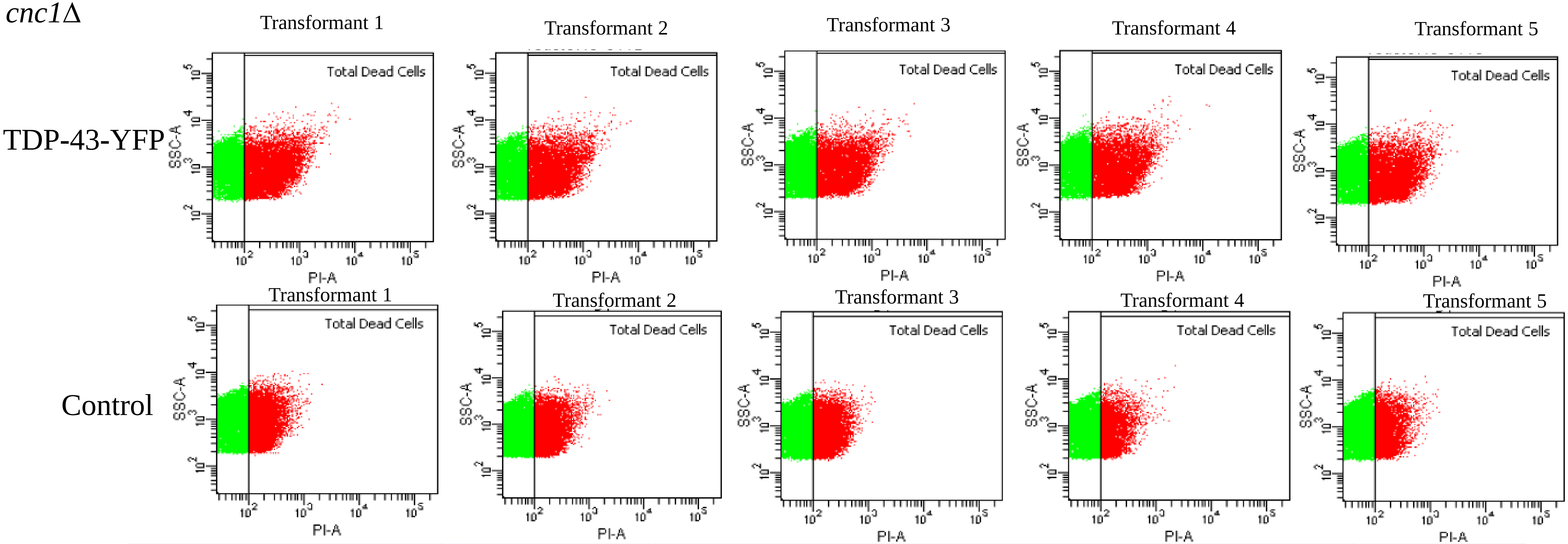

| | | Transformant 1 | Transformant 2 | Transformant 3 | Transformant 4 | Transformant 5 |
|---|---|---|---|---|---|---|
| | Total cells | 50000 | 50000 | 50000 | 50000 | 50000 |
| | Dead cells | 8747 | 8048 | 7819 | 8181 | 7815 |
| *cnc1Δ* TDP-43-YFP | % of Dead cells | 17.494 | 16.096 | 15.638 | 16.362 | 15.63 |
| | | Transformant 1 | Transformant 2 | Transformant 3 | Transformant 4 | Transformant 5 |
| | Total cells | 50000 | 50000 | 50000 | 50000 | 50000 |
| | Dead cells | 9689 | 11018 | 9896 | 6073 | 6208 |
| *cnc1Δ* Control | % of Dead cells | 19.378 | 22.036 | 19.792 | 12.146 | 12.416 |

**Supplementary Figure 2S**

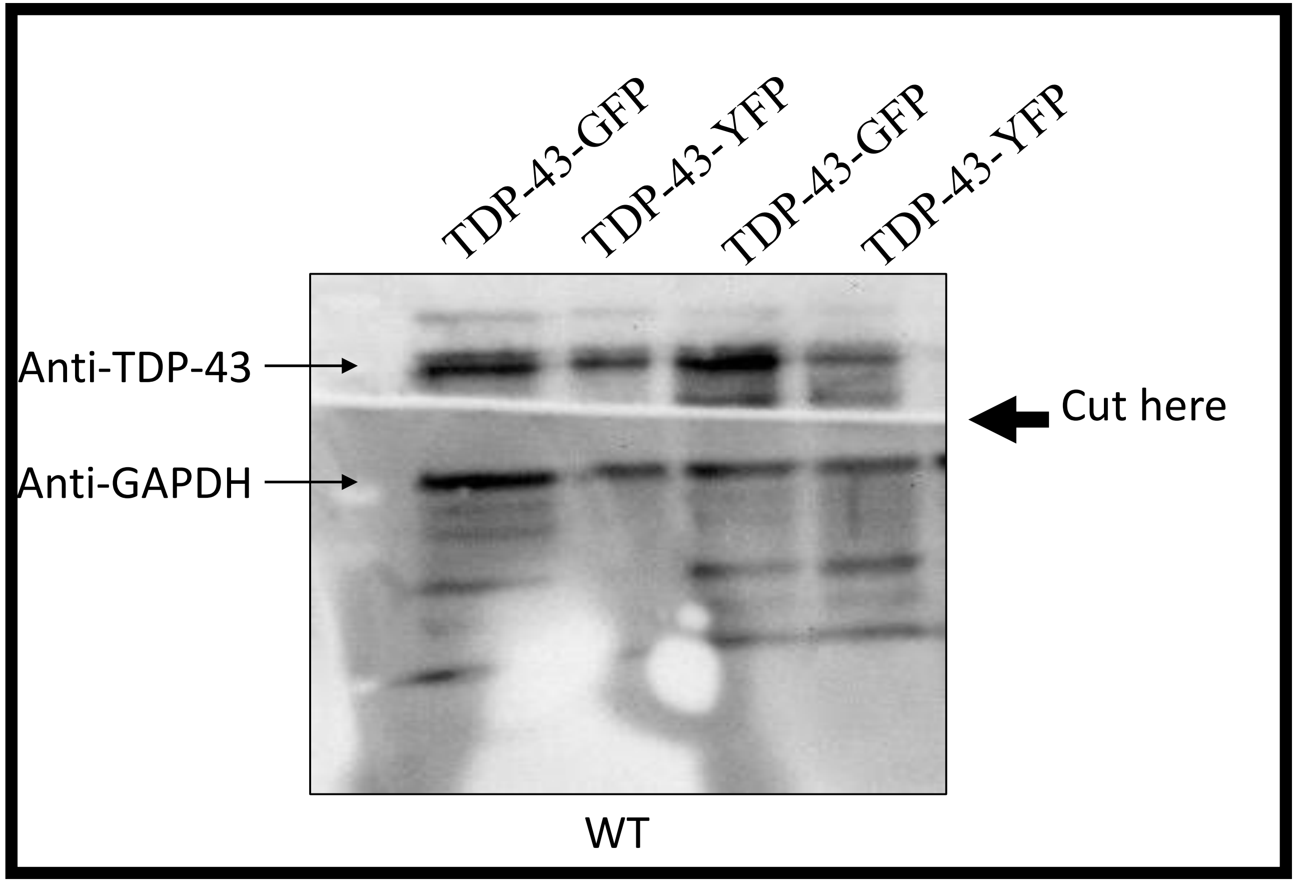

**Supplementary Figure 2S**: **TDP-43 expression from high and low copy number plasmid estimated by western blotting**

Two fusion proteins of TDP-43 namely TDP-43-YFP expressed from a low copy number plasmid or TDP-43-GFP expressed from a high copy number plasmid, both under *GAL1* promoter control, were checked for relative TDP-43 protein levels by western blotting using anti-TDP-43 antibody after 24 hours of expression. Endogenously expressed GAPDH protein was simultaneously probed with anti-GAPDH antibody, as the protein loading control. TDP-43-GFP exhibited many-fold higher expression than TDP-43-YFP.

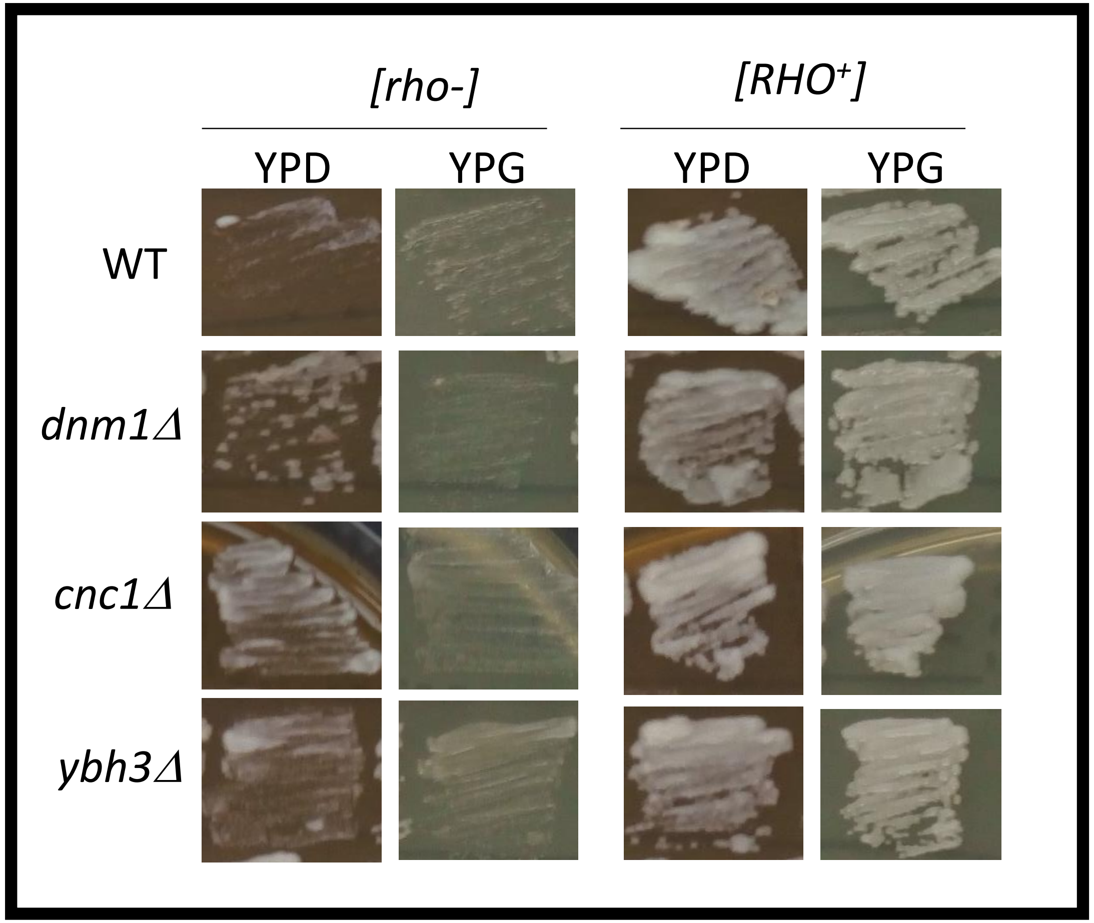

**Supplementary Figure 3S: Generation of yeast petite strains *[rho⁻]***

Isogeneic petite [*rho*⁻] versions of wild-type (WT), *dnm1Δ*, *ybh3Δ* and *cnc1Δ* yeast strains were created by growing the [*RHO*⁺] strains on YPD supplemented with 0.5 mg/ml ethidium bromide (EtBr). Cells from the EtBr plates were streaked on to YPD to obtain single colonies. Several colonies from YPD were analysed for the respiration deficient mutants (i.e. conversion to [*rho*⁻] or petite strain) by checking for their lack of growth on the complex media containing 2% glycerol (non-fermentable) as the only carbon source (YPG).

**Supplementary Figure 4S**

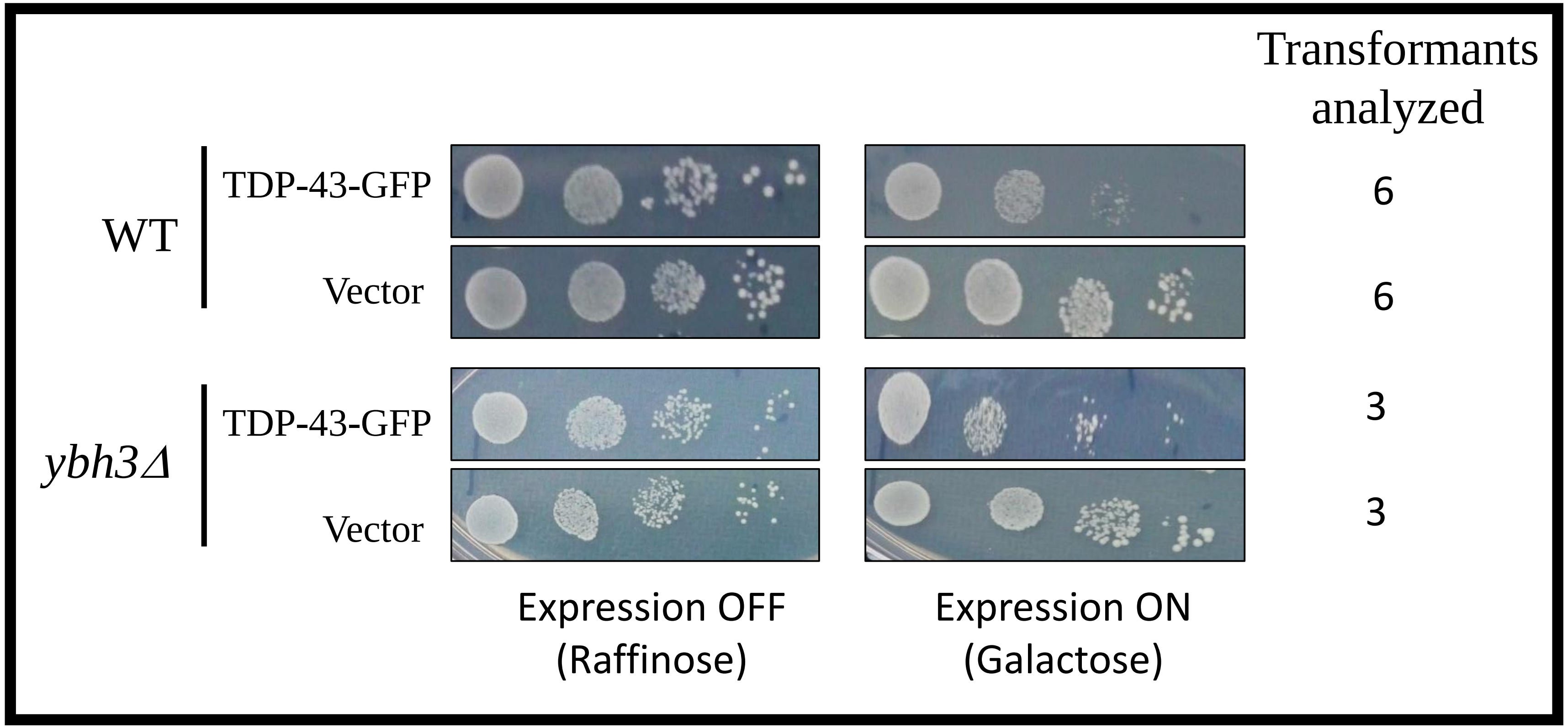

**Supplementary Figure 4S: Rescue of toxicity of highly over-expressed TDP-43-GFP in *ybh3Δ* yeast**

Wild-type (WT) or *ybh3Δ* yeast were transformed with high copy number, *GAL1* promoter-driven plasmid encoding TDP-43-GFP or the empty vector. Serial dilution (10-fold) growth assays were performed onto galactose-containing media where the TDP-43-GFP expression is turned on and also for the cell number controls onto raffinose-containing plates where the *GAL1* promoter is switched-off. Serial dilution (10-fold) growth assay of *ybh3Δ* transformed with a *GAL1*-driven high copy number plasmid encoding TDP-43-GFP showed that TDP-43-GFP toxicity was partially rescued in comparison with the WT TDP-43-GFP.

**Supplementary Figure 5S: Oxidative stress-induced cytoplasmic localization of Cyclin C upon TDP-43 co-expression**

Cyclin C-YFP and TDP-43-DsRed, both under *GAL1* promoter, were expressed with 2% galactose for 24 hours in the wild-type BY4742 yeast. Yeast cells were then stained with Hoechst stain for visualising the cellular nuclei. In the absence of TDP-43 expression, Cyclin C-YFP exhibited mostly nuclear localization and upon TDP-43 co-expression, Cyclin C-YFP significantly moved into the cytoplasm. When the anti-oxidant, N-acetyl-cysteine (NAC) was added along with the cells expressing both Cyclin C-YFP and TDP-43-DsRed, the cytoplasmic translocation of Cyclin C-YFP was significantly reduced. Representative images from each transformant are givrn below. Images were acquired using DM2500 Leica Fluorescence microscope.

#### Supplementary Figure 5S

Cyclin C-YFP + Vector

Transformant 1

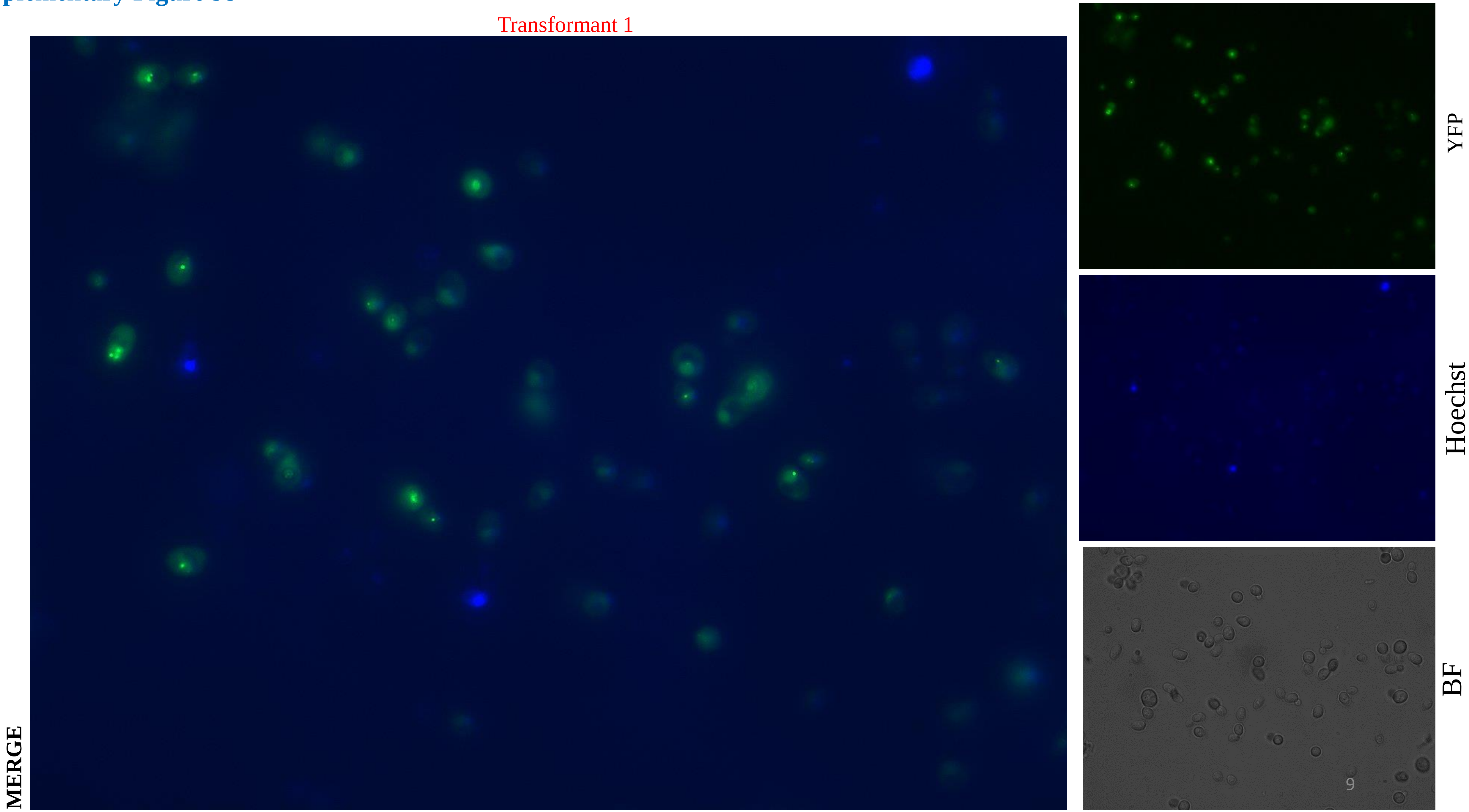

#### Supplementary Figure 5S

Cyclin C-YFP + Vector

Transformant 2

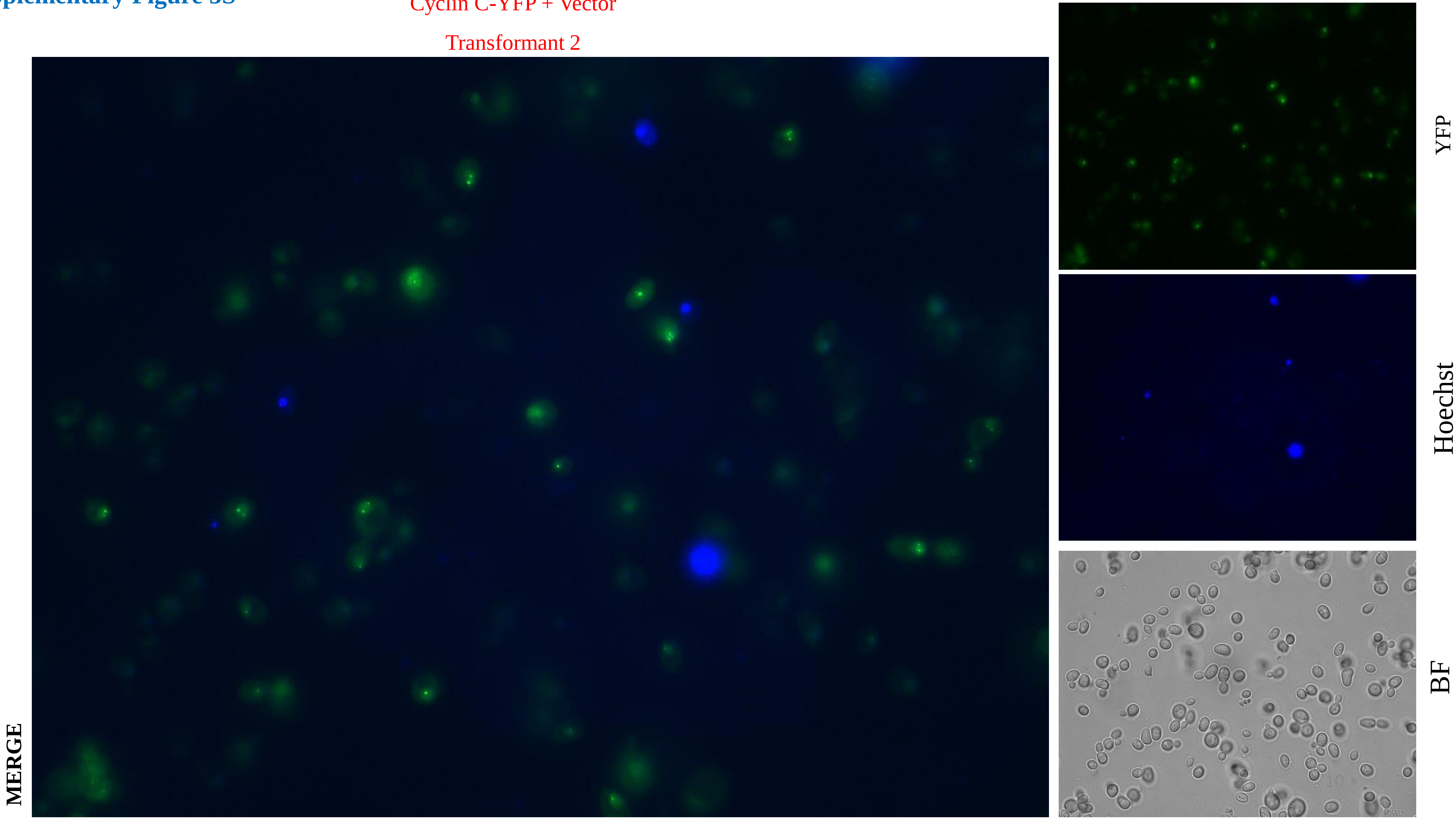

#### Supplementary Figure 5S

Cyclin C-YFP + Vector

Transformant 3

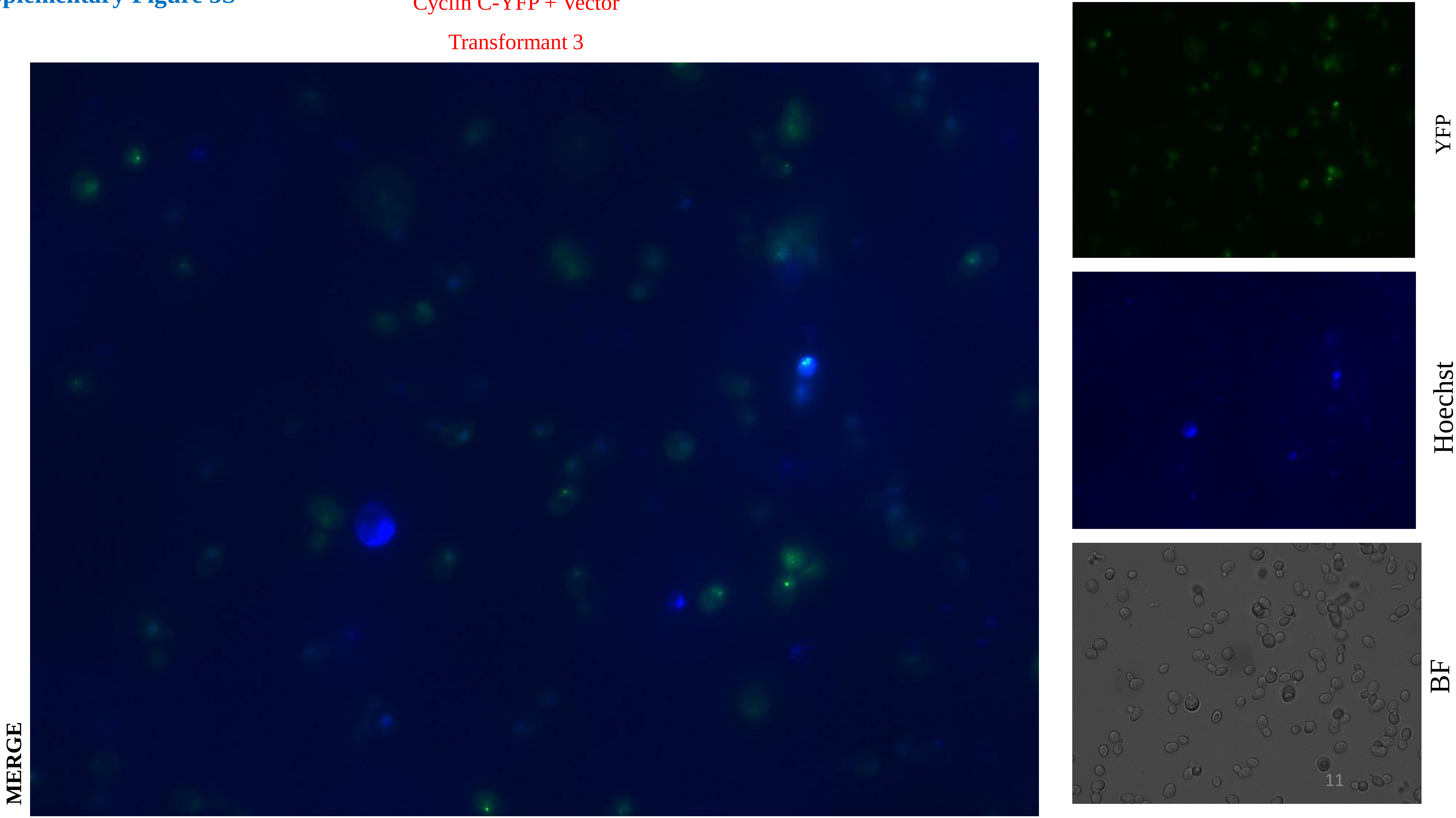

#### Supplementary Figure 5S

Cyclin C-YFP + TDP-43-DsRed

Transformant 1

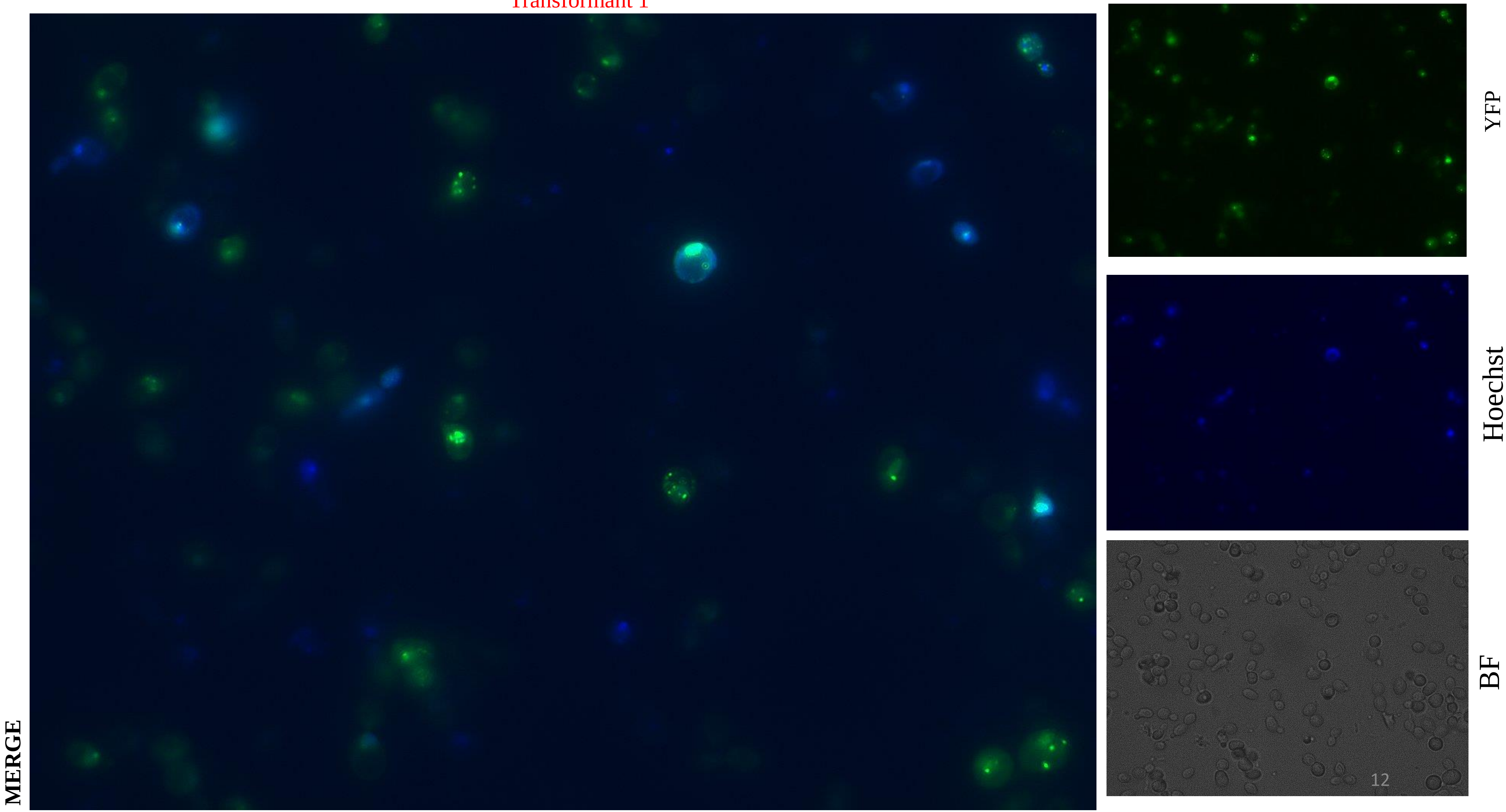

#### Supplementary Figure 5S

Cyclin C-YFP + TDP-43-DsRed

Transformant 2

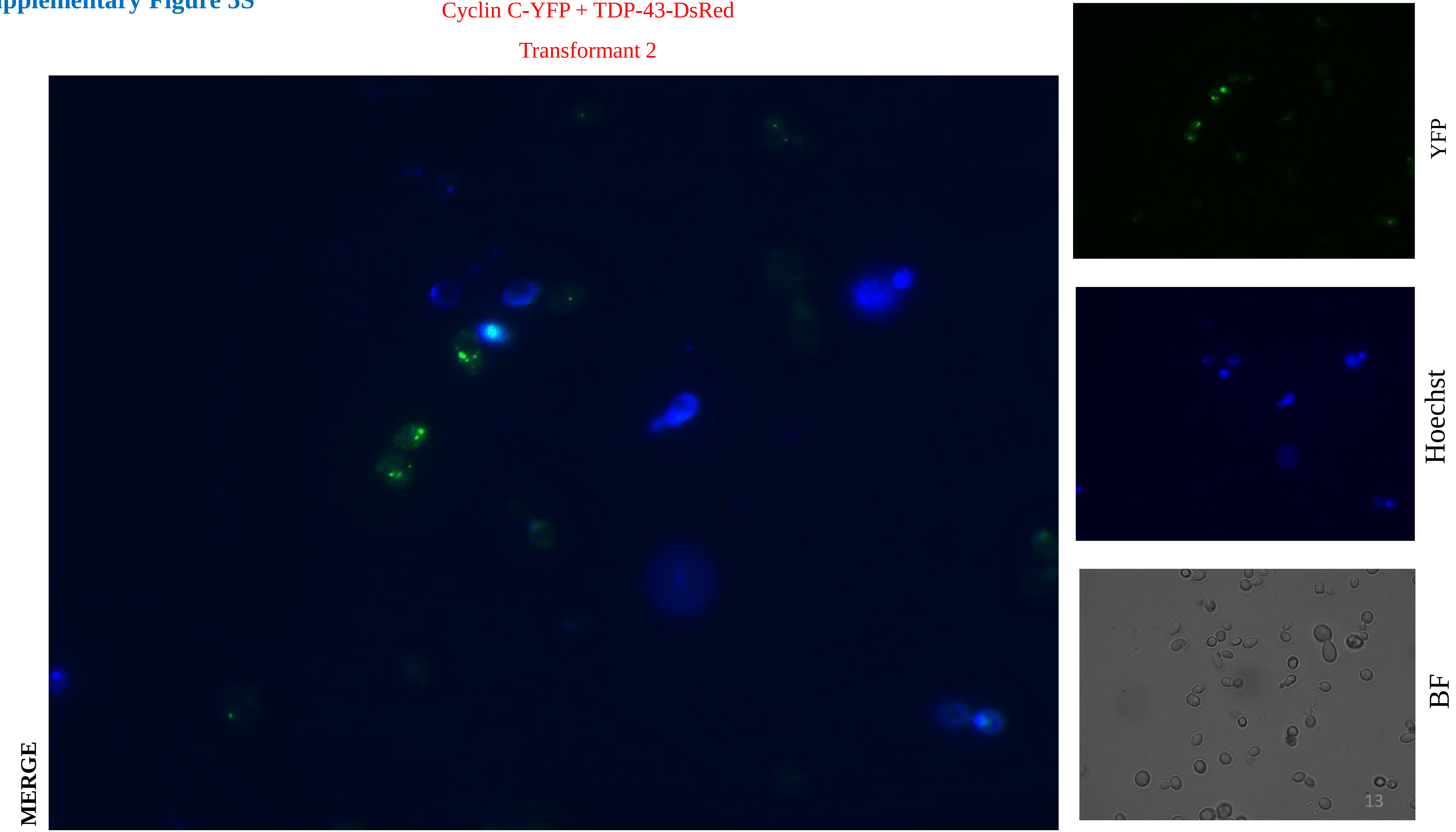

**Supplementary Figure 5S**

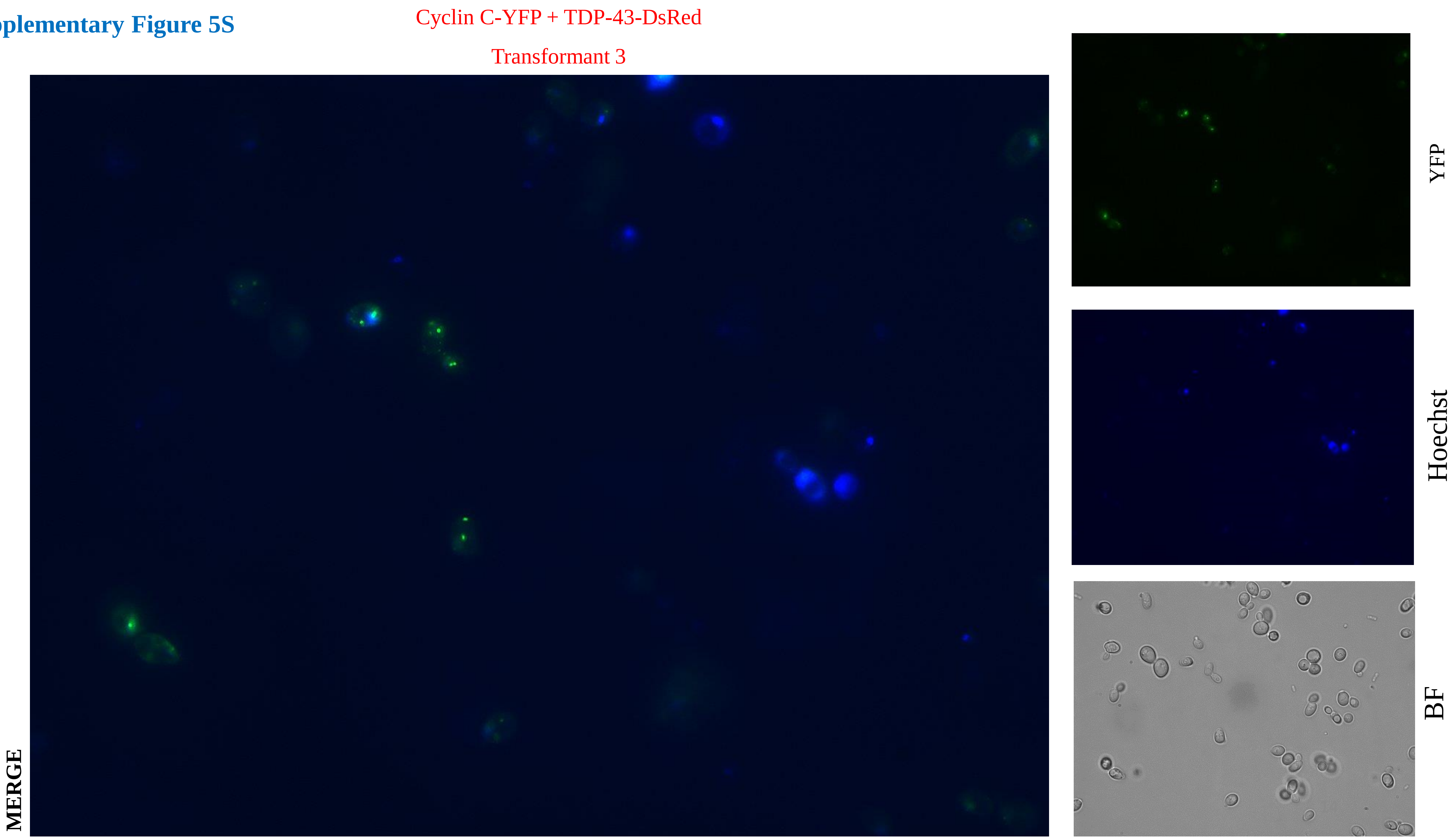

### Supplementary Figure 5S

Cyclin C-YFP + TDP-43-DsRed + 5 mM NAC

Transformant 1

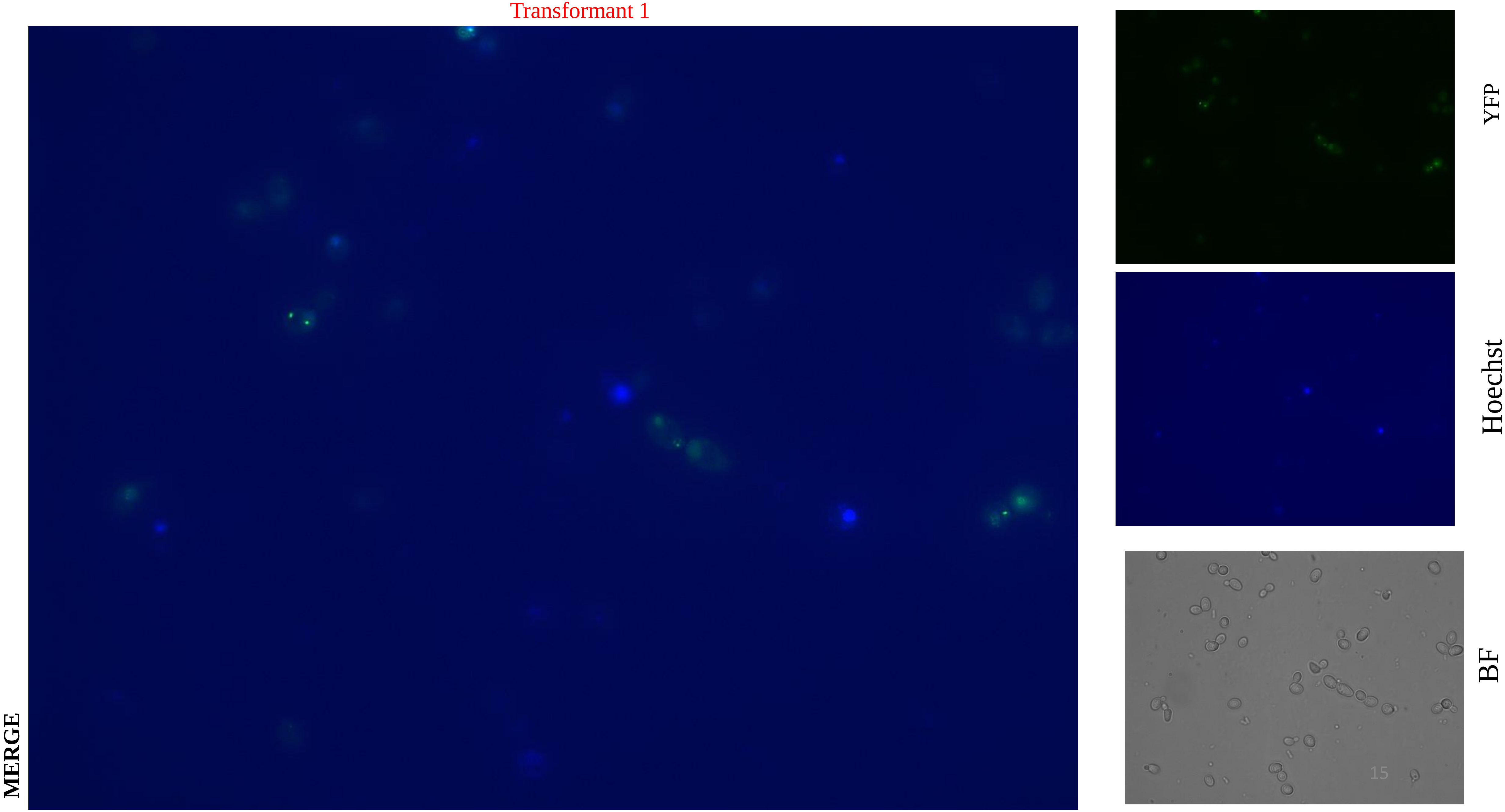

**Supplementary Figure 5S**

Cyclin C-YFP + TDP-43-DsRed + 5 mM NAC

Transformant 2

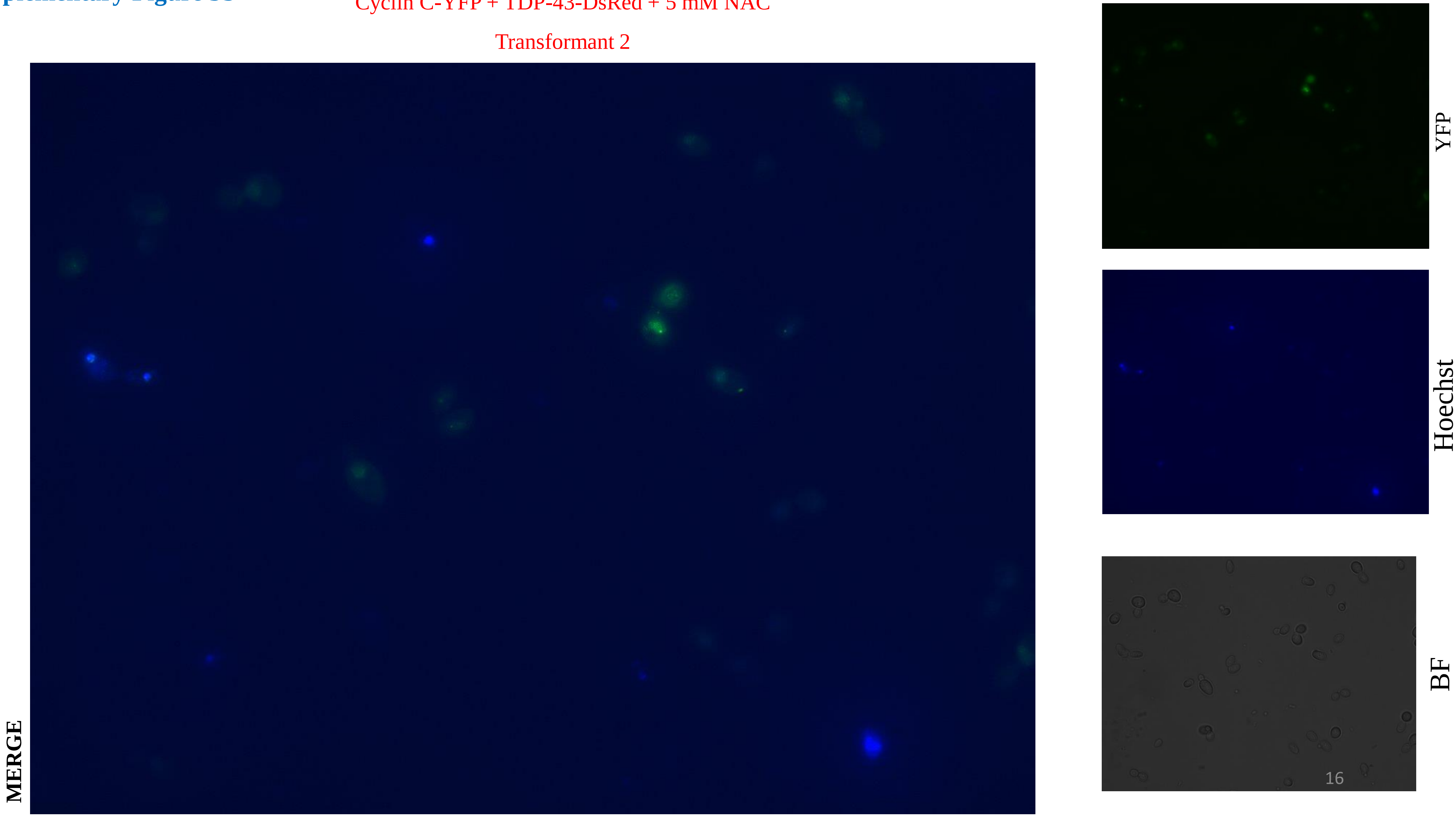

#### Supplementary Figure 5S

Cyclin C-YFP + TDP-43-DsRed + 5 mM NAC

Transformant 3

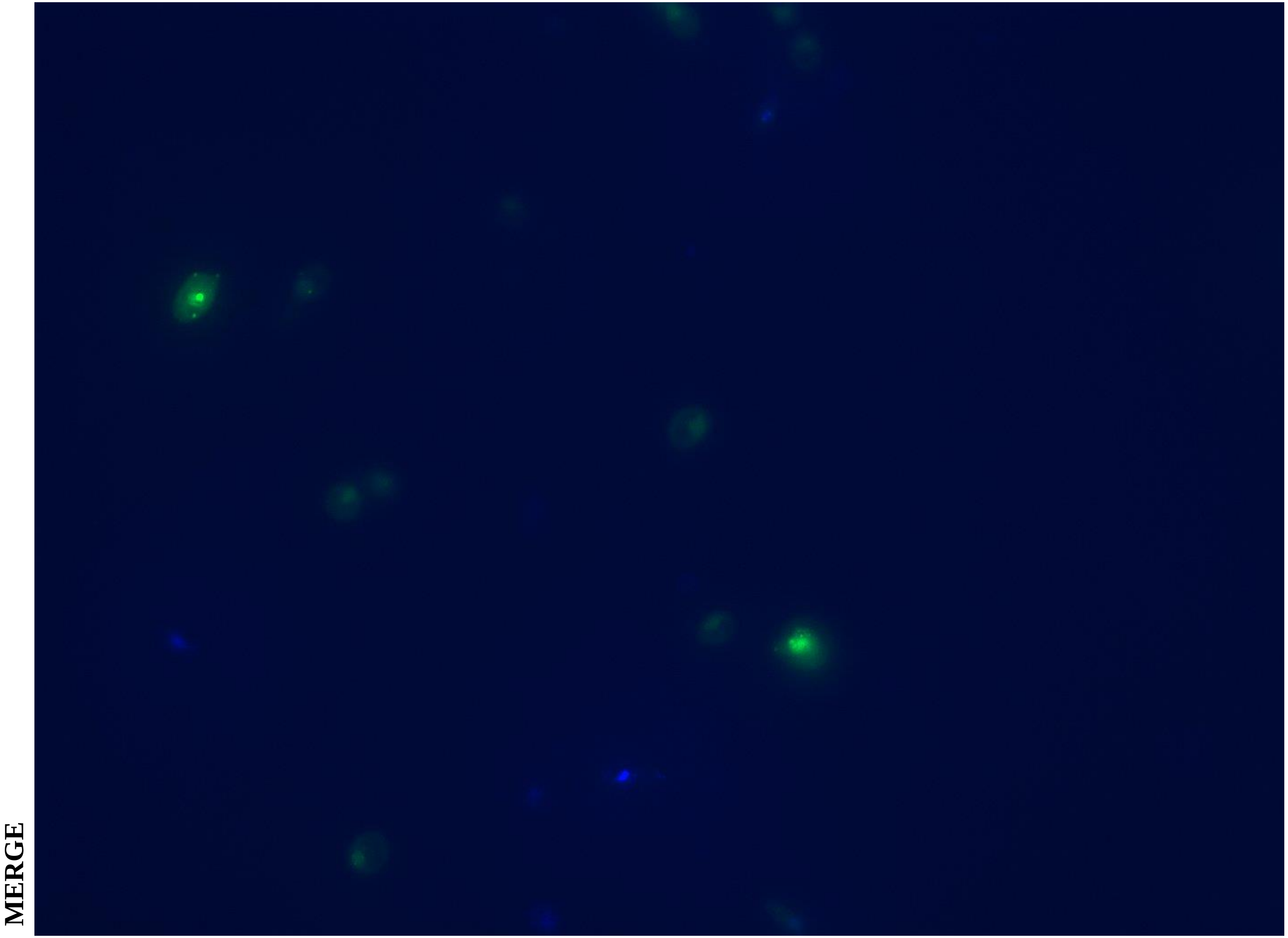

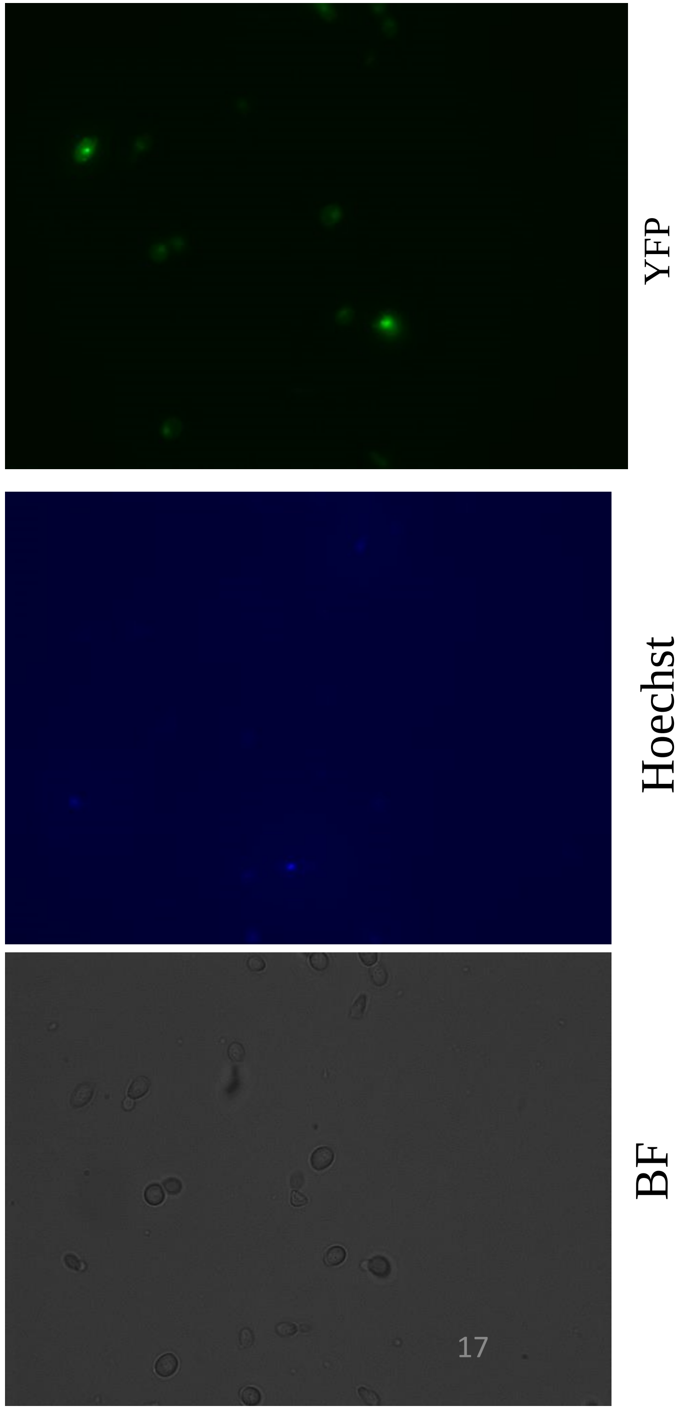

### Supplementary Figure 6S

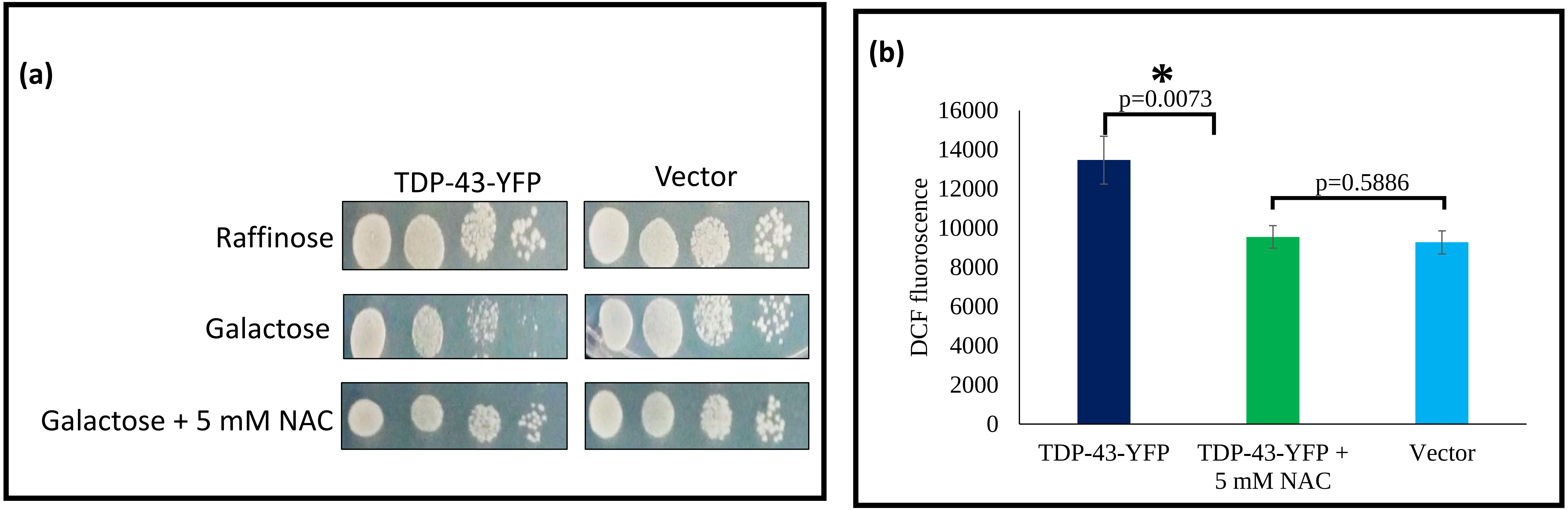

**Supplementary Figure 6S**: **Modulation of TDP-43 induced cytotoxicity and oxidative stress by N-acetyl-cysteine**

**a.** Wild-type (WT) yeast was transformed with low copy number, *GAL1* promoter-driven plasmids encoding TDP-43-YFP or the empty vector. Serial dilution (10-fold) growth assays were performed onto galactose-containing media where the TDP-43-YFP expression is turned-on and also for the cell number controls onto raffinose-containing plates where the *GAL1* promoter is switched-off. Toxicity of TDP-43 expression was also simultaneously checked on a galactose-containing media containing plate supplemented with 5 mM of N-acetyle cysteine (NAC). It was found that TDP-43 cytotoxicity was relieved in the presence of NAC.

**Supplementary Figure 6S: Modulation of TDP-43 induced cytotoxicity and oxidative stress by N-acetyl-cysteine**

**b**. 2’,7’ Dichlorofluoroscein diaceate (DCFDA) assay was performed to assess the levels of reactive oxygen species (ROS) in the yeast cells expressing TDP-43-YFP or vector as described previously (Bharathi et al, Yeast, 33, 607-620, 2016)). For this, the yeast cultures were grown overnight in SRaf-Ura and inoculated into fresh SGal-Ura for the overexpression of the TDP-43-YFP. Yeast cells harbouring an empty vector (pRS416) were induced and grown similarly and included as a control for comparison of the ROS levels. DCFDA assay was performed as described earlier (Bharathi et al, Yeast, 33, 607-620, 2016). DCF fluorescence emission intensity at 524 nm was recorded using Enspire multimode microplate reader (Perkin Elmer, USA) after excitation at 485 nm. Expression of TDP-43 in the yeast elicited elevated oxidative stress which was significantly lowered with the addition of N-acetyl-cysteine. Data represents the Reactive oxygen species (ROS) levels from three independent transformants and the error bars represent standard deviation. p-values were determined with un-paired t-test. * in the p-value represents statistical significance.
